# Ligand dependent gene regulation by transient ERα clustered enhancers

**DOI:** 10.1101/541540

**Authors:** Bharath Saravanan, Deepanshu Soota, Zubairul Islam, Ranveer Jayani, Rajat Mann, Umer Farooq, Sweety Meel, Kaivalya Walavalkar, Srimonta Gayen, Anurag Kumar Singh, Sridhar Hannenhalli, Dimple Notani

**Affiliations:** National Centre for Biological Sciences, Tata Institute of Fundamental research, Bangalore, India, 560065; School of Medicine, University of California, La Jolla, 92093; Molecular Reproduction, Development and Genetics, Indian Institute of Science, Bangalore, India, 560012; Center for Bioinformatics and Computational Biology, University of Maryland, College Park, MD, 20742; Sastra Deemed University, Thanjavur, Tamil Nadu, India, 613401; The University of Trans-disciplinary Health Sciences and Technology, Bangalore, India, 560064

## Abstract

Unliganded nuclear receptors have been implicated in ligand-dependent gene regulation. However, the underlying mechanisms are not fully understood. Here we demonstrate that unliganded ERα binds to specific sites in the genome thereby pre-marking them as future functional enhancers. Upon ligand exposure, ERα binds to several EREs relatively proximal to the pre-marked, or persistent, ERα-bound sites. Interestingly, the persistent sites interact extensively, via chromatin looping, with the proximal transiently bound sites forming ERα clustered enhancers in 3D. CRISPR-based deletion of *TFF1* persistent site disrupts the formation of its clustered enhancer resulting in the loss of E2-dependent induced expression of TFF1 and its neighboring genes within the same cluster. The clustered enhancers overlap with nuclear ERα puncta that coalesce in a ligand-dependent manner. Furthermore, formation of clustered enhancers, as well as puncta, coincide with the active phase of signaling and their later disappearance results in the loss of gene expression even though persistent sites remain bound by ERα. Our results establish the role of persistent unliganded ERα binding in priming enhancer clusters in 3D that drive transient, but robust, gene expression in a ligand-dependent fashion.

## Introduction

Estrogen receptor alpha (ERα) translocates into the nucleus upon ligand binding where it regulates gene expression by binding to distal regulatory elements (Li et al., 2013, Liu et al., 2014, Hah et al., 2011). Interestingly, a basal quantity of ERα is found in the nucleus even in the absence of ligand. Unliganded nuclear ERα binding was recently shown to mark future functional enhancers, whereby these pre-bound ERα sites serve as seeds, around which multiple ERα-bound peaks emerge after ligand exposure (Bojcsuk et al., 2017, Caizzi et al., 2014). However, mechanisms underlying these intriguing observations, and their role in gene regulation, are not entirely clear. For instance, it is not known whether unliganded pre-bound ERα sites are absolutely required for such ligand-dependent ERα-clustering at the genomic level and the downstream gene regulation. The estradiol (E2)-dependent gene expression peaks at 1 hour post stimulation and drops at 3 hours (Hah et al., 2011). However, whether the transient response to signaling is driven by transient existence of enhancers is not known. Furthermore, whether the constituent enhancers within clustered enhancers exhibit E2-dependent physical proximity to form a functional unit is yet to be investigated.

Super-enhancers exhibit high density of coactivators caused by presence of multimeric enhancer units. Intrinsically disordered regions (IDR) in these coactivators mediate their molecular condensation on super enhancers (Sabari et al., 2018). ERα, with its co-activator Med1, was recently shown to form such condensates (Boija et al., 2018). However, whether *bona fide* ERα clustered enhancers are tethered in such ERα condensates is not known. Importantly, whether enhancer condensates continue to exist during the entire interval when the ligand is present, or whether they exist only during the active signaling phase marked by robust transcription is not known.

Here we show that upon E2 treatment, as expected, ERα binds to numerous locations across the genome. As recently observed (Bojcsuk et al., 2017), the new ligand-dependent sites are significantly organized as clusters in relative proximity to pre-existing, or *persistent*, ligand-independent ERα-bound sites. However, we find that the ligand-dependent enhancer clusters (LDEC) are distinct from previously reported super-enhancers (Hnisz et al., 2013). LDECs exhibit extensive chromatin looping among constituent ERα sites within as well as across LDECs in 3D. Specifically, LDECs emerge robustly and transiently upon E2 treatment and their disappearance at 3 hours post-treatment coincides with the loss of eRNA at constituent enhancers and their cognate target gene expression as well as drop in ERα protein levels. LDECs may include multiple genes whose expressions are concomitant with LDEC existence We show via CRISPR deletion that persistent sites are absolutely required for ligand-induced binding of ERα to the neighboring EREs. ImmunoFISH experiments indicate that these 3D ERα clusters overlap with ERα puncta only after ligand stimulation. Further, ERα forms condensates on ERα-clustered enhancers by coalescence upon E2 treatment.

Overall, our results establish and clarify the role of unliganded ERα binding in priming enhancer clusters in 3D that drive transient, but robust, gene expression in a ligand-dependent fashion. Our work suggests a model of E2-induced gene regulation where during active phase, liganded ERα decorates the EREs closer to pre-marked unliganded ERα site. These pre-marked and new ERα-bound sites exhibit E2-dependent extensive chromatin looping forming functional LDEC that drives gene expression. These LDECs correspond to ERα puncta in the nucleus that emerge upon E2 stimulation. Finally, upon ERα degradation at 3h or upon deletion of persistent site, these clusters disappear leaving persistent site behind still bound by ERα as bookmark, but that alone is unable to drive gene expression.

## Results

### ERα binds in clusters around pre-existing ERα-bound sites upon ligand stimulation

ERα binds predominantly in the intergenic regions of the genome upon E2 stimulation (Fig S1A, B and C). Further, we tested if ERα bound sites exhibit genomic clustering as has been reported recently (Bojcsuk et al., 2017). Consecutive ERα peaks within 20kb were considered to be part of the same genomic cluster. We identified 1514 ERα clusters containing at least three ERα peaks in each cluster (see Methods). As a control, we repeated the clustering analysis based on hundred iterations of randomly selected 21,834 ERE motif, or DHS (Methods), and found that 7% of all EREs (482 clusters) and 22% of the DHS sites (1304 clusters) clustered compared with 30% of total ERα peaks (Fig 1A). These results strongly support the tendency of ERα peaks to cluster on the genome. To investigate if the existence of clusters is dependent on ligand stimulation, we compared the post-E2 ERα-bound peaks with those in non-stimulated MCF7 cells. We observed 6659 peaks in the untreated cells, of which 3779 peaks were also found in the post-E2 condition (Fig 1B); we refer to such sites as *persistent* sites and the newly acquired 18,055 E2-dependent sites as *transient* sites. Further, we found that a majority of the above 1514 clusters had at least one persistent site, suggesting their potential role in seeding the formation of genomic clusters of ERα sites upon E2 treatment. Further, contrasting persistent and transient sites, we found persistent sites to have higher affinity of ERα binding relative to transient sites (Fig 1C). We binned persistent sites into three quantiles based on ERα binding strength post-E2. Persistent sites in general, and the 3^rd^ quantile persistent sites in particular, exhibited stronger ERα binding as well as significantly greater DHS in both post and pre-E2 treated cells (Fig 1C; Methods). We further probed the binding strength differences between persistent and transient sites via motif enrichment analyses. Not surprisingly, both the persistent and the transient sites were highly enriched for EREs (Fig. 1D). Interestingly, however, the only motifs enriched uniquely in the persistent sites were for FOX proteins where FOXA1 and FOXA2 motifs had the most significant p-values (Fig. 1D). FOXA1 is a known pioneer *cis* co-factor for ERα binding (Swinstead et al., 2016, Hartado et al., 2011, Carroll et al., 2005) and is frequently mutated in breast tumors (Fu et al., 2016). We confirmed the exclusive preference of FOXA1 binding on persistent sites (Fig S1D). These results indicate that FOXA1 bound sites along with classical ERE motifs are the hallmarks of persistent sites which may give rise to the clustered binding of ERα around these persistent sites on signaling. We observed that most of the robust E2-induced genes are harbored around these clusters for example *CCND1, NRIP1, TFF1* and *GREB1* (Fig 1E). These clusters are marked by H3K27ac suggesting their enhancer activity (Fig 1E). Finally, relative to transient sites, persistent sites seem to be under a greater purifying selection as evidenced by higher cross-species conservation (Phast-cons scores, Methods) (Fig S1E). Furthermore, the ERα peaks within clusters exhibited an intriguing pattern, with the strongest binding at persistent sites showing a gradual decrease in ERα binding strength at consecutive ERα sites as we go farther from the persistent site in either direction (Fig S2A). However, DHS signal on these transient sites remained unchanged (Fig S2B), suggesting a gradually decreasing sphere of influence of the persistent site.

**Figure 1.**
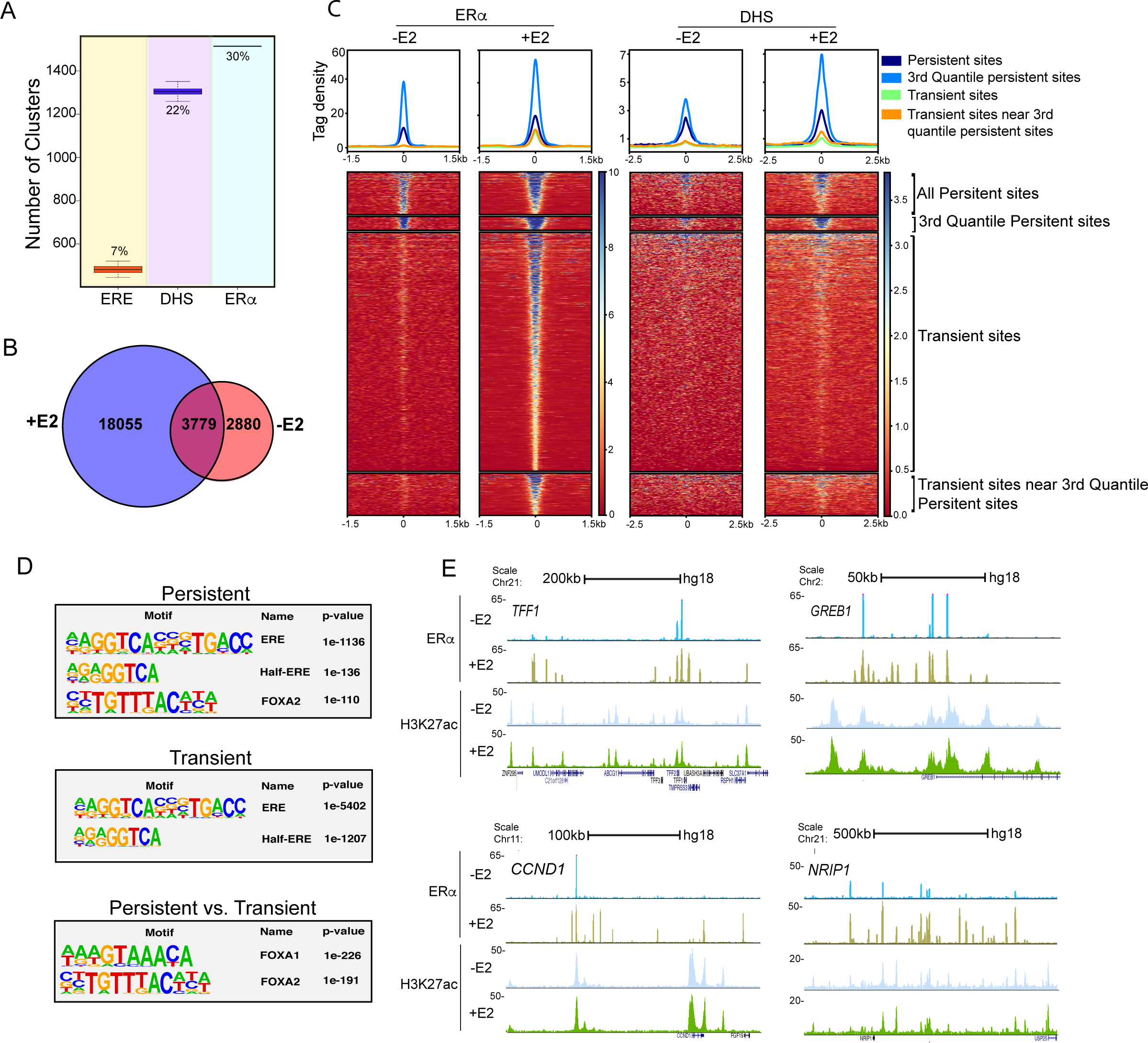
Estrogen receptor binds in clusters around pre-existing ERα-bound sites upon ligand stimulation. (A) ERα shows highest propensity for genomic clustering compared to ERE and DHS. Y-axis represents the number of clusters with at least 3 sites (ERα, ERE, or DHS individually) with less than 20kb between consecutive sites. (B) Venn diagram showing 3779 persistent ERα peaks being common in control and E2 treated cells. (C) Heat map exhibiting binding strength of ERα (left panel), and DHS (right panel) on all persistent, strongest persistent (3^rd^ quantile), all transient, and transient sites clustered around a persistent site. (D) (top panel) Known Motif enrichment analysis identifies full ERE in both persistent (p=10^-1136^) and transient sites (p=10^-5402^) (Middle Panel) whereas FOXA1 is enriched uniquely in persistent with p=10^-226^ (Top and Lower Panels). (E) UCSC genome browser snapshots showing the binding of ERα and H3K27ac status on ERα clustered enhancers for four of the robustly E2-induced target genes – *TFF1*, *GREB1*, *NRIP1*, and *CCND1* in unliganded and in E2 conditions. The boxplots depict the minimum (Q1-1.5*IQR), first quartile, median, third quartile and maximum (Q3+1.5*IQR) without outliers.

### ERα enhancer clusters but not the conventional super-enhancers regulate E2-dependent genes

Our detected ERα enhancer clusters share salient properties with super-enhancers, namely, genomically clustered constituent enhancers (Hnisz et al., 2013), raising the possibility that ERα clustered enhancers correspond to super-enhancers. Using the Rose tool (Hnisz et al., 2013) to detect super-enhancers, in the post-E2 cells, 858 H3K27ac-based super-enhancers and 390 ERα binding strength-based super-enhancers were identified in E2 condition. We first confirmed that all 390 ERα super-enhancers (LDEC) overlapped with our detected 1514 clusters(Fig 1). However, only 79 of the ERα super-enhancers overlapped with H3k27ac super-enhancers (Fig. 2A). Although, as expected, H3K27ac super-enhancers exhibited higher enrichment of H3K27ac marks as compared to ERα super-enhancers (Fig 2C), importantly, upon E2 stimulation H3K27ac and ERα occupancy did not change on H3K27ac super-enhancers, whereas ERα occupancy but not the H3K27ac increased significantly on ERα super-enhancers (Fig. 2B and C). Interestingly, DHS signal was strengthened on both classes of super-enhancers (Fig 2D) suggesting that modulation of H3K27ac super-enhancers by E2 is independent of direct ERα binding. Super-enhancers have been shown to activate their target genes robustly compared to typical enhancers. To compare the strength of gene activation by the two classes of super-enhancers, we compared the normalized GRO-seq (Hah et al., 2011) tag counts of 4 closest genes to each enhancer in the two classes before and after E2 induction. We observed significant E2-dependent upregulation of genes closer to ERα super-enhancers compared to the H3K27ac super-enhancers, suggesting that ERα super-enhancers but not the H3K27ac super-enhancers robustly activate their target genes in an E2-dependent manner (Fig 2E and F).

**Figure 2.**
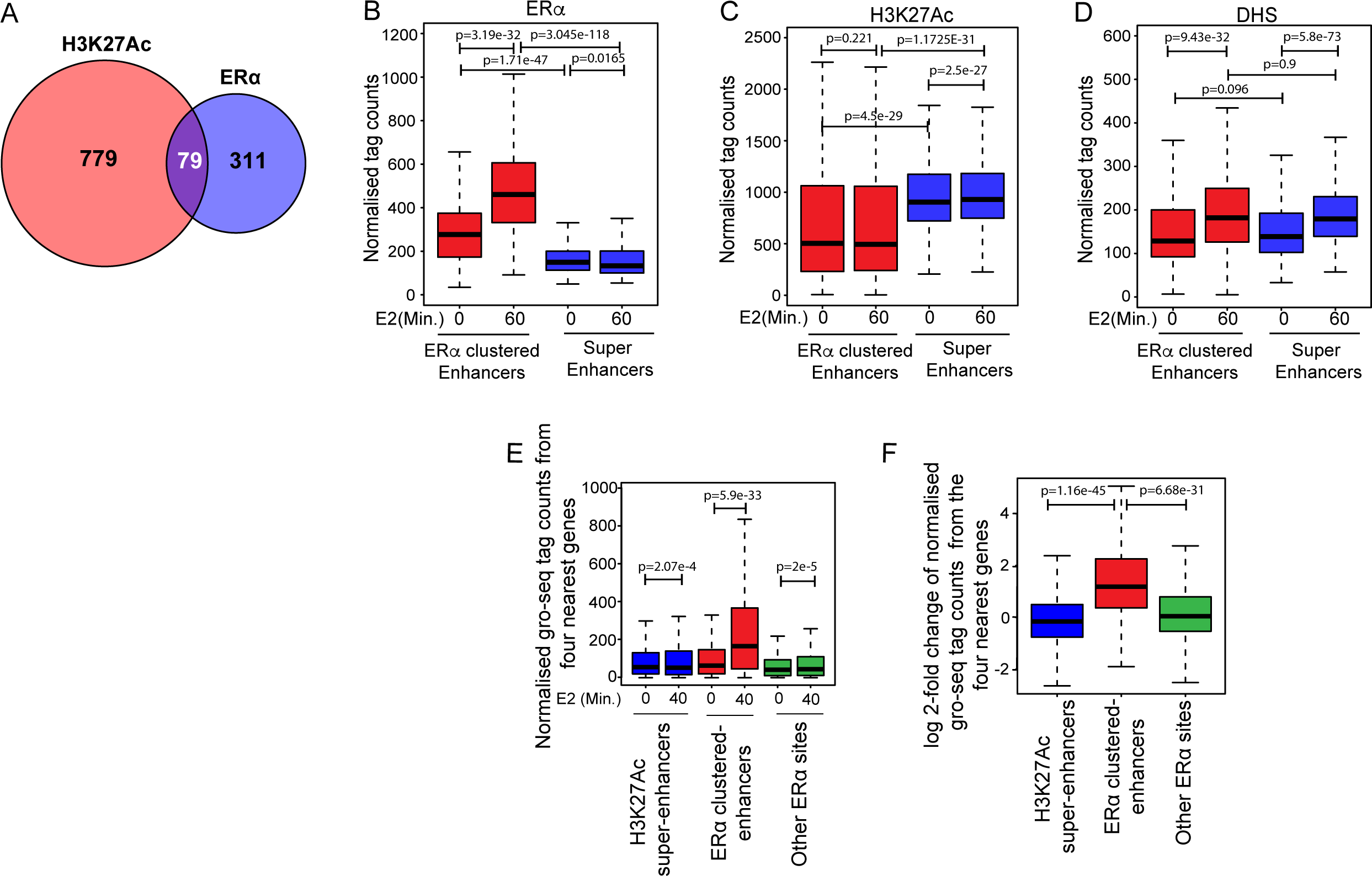
ERα super-enhancers but not conventional H3K27ac super-enhancers control E2 target genes. (A) Venn diagram showing overlap of 79, H3K27ac super-enhancers with ERα super-enhancers. (B-D) ERα binding strength (B), H3K27ac enrichment (C), and DHS signal (D) on H3K27ac super-enhancers and ERα super-enhancers in control and 40’ post-E2 treated cells show higher levels of ERα induction on ERα super-enhancers, higher H3K27ac enrichment on H3K27ac super-enhancers, but similar DHS induction in both categories of enhancers. (E) Expression of genes closer to H3K27ac super-enhancers and other ERα sites not in cluster do not change upon E2 treatment but the genes that are closer to ERα super-enhancers are highly induced upon E2 treatments. (F) Pre-E2 to post-E2 log2 fold changes in expression of genes closer to different categories of enhancers as mentioned in panel C. All p-values were calculated by either Wilcoxon rank sum test or Wilcoxon signed rank test. The boxplots depict the minimum, first quartile, median, third quartile and maximum without outliers.

### Genomically clustered ERα sites exhibit 3D proximity with each other and with target gene promoter(s)

Genes that are induced in a ligand-dependent manner exhibit induced physical proximity to their enhancers in a ligand-dependent manner (Li et al., 2013, Hsieh et al., 2014, Li et al., 2016). Given our observation that the genes near LDECs exhibit robust E2-dependent upregulation, we hypothesized that for the LDECs to act as a regulatory unit in a way similar to the conventional super-enhancers, the constituent persistent and transient ERα sites within a LDEC should physically interact with each other and also with the target gene promoter. Since persistent ERα sites exhibit greater levels of ERα binding, we tested if these sites show relatively higher degree of physical interactions with other ERα-bound sites. Using publicly available ChIA-PET data on ERα in MCF7 cells (Fullwood et al., 2009), we found that overall the persistent sites exhibit a higher degree of interactions compared to the transient sites, and this tendency is even greater on highest occupancy persistent sites (3^rd^ quantile) (Fig 3A). Further, to assess whether LDECs are contained within a single Topologically Associated Domains (TAD) or span over multiple TADs, we interrogated HiC-inferred TADs from E2-treated MCF7 cells (Rodriguez et al., 2018). We found that 70% of LDECs are within a TAD, which indicates topological constrains on LDECs. Further, including HiC data from untreated cells (Rodriguez et al., 2018) we noted that TADs containing LDEC exhibit a strengthening of intra-TAD over inter-TAD interactions upon E2 stimulation (Fig. 3B). These observations suggest potentially strengthened looping between persistent and transient sites, and with the target gene promoters, within the LDEC upon ligand addition. To capture such interactions, we investigated in depth specific LDECs using the available ERα ChIA-PET data. Interestingly, for the LDEC in the TFF1 gene locus, the persistent and its neighboring transient sites exhibited strong interactions with each other and with TFF1 promoter (Fig 3C). Interrogating longer-range interactions, we found interactions between TFF1 LDEC and another LDEC near *UMODL1* gene 200 kb away (Fig 3C), indicating the presence of extensive looping within LDEC and also between neighboring LDECs. Intra-LDEC and LDEC-promoter interactions were also observed for other tested genes such as *NRIP1* (Fig 3D) and *GREB1* (Fig S3A). Because ChIA-PET was performed in presence of E2, we next aimed to determine whether the observed interactions were E2-dependent. Due to relative genomic proximity of ERα sites within a cluster, requiring greater sensitivity in spatial proximity determination, we performed 5C on all clustered enhancers on Chr21, as it harbors the most robustly E2-induced genes, viz., *TFF1*, *NRIP1*, and the lncRNA *DSCAM-as*. These genes span the entire length of the long arm of acrocentric Chr21. The 5C library contained forward and reverse oligos for all active enhancers (Liu et al., 2014) and the respective promoters. Oligos derived from ENCODE desert region on Chr16 were used to normalize the digestion and ligation efficiency biases between the libraries as reported earlier (Sanyal et al., 2012). The 5C was performed in ICI and 1hr E2-treated MCF-7 cells, number of reads and contacts are mentioned in (Table S1). The data in E2 treated and untreated cells clearly showed an overall comparable 5C matrix ruling out any bias. Interestingly, the overall normalized 5C data showed a strong diagonal upon E2 induction reflecting enriched cis-interactions (Fig 3E). Similar enrichment in cis-interactions upon E2 treatments has been reported recently (Rodriguez et al., 2018). To interrogate the observed E2-dependent strengthening interactions within LDECs (Fig 3B), we plotted the Chr21 normalized reads arising from the binned reads overlapping genomic regions around and within the cluster (Method and TableS2). A snapshot of *TFF1* and *NRIP1* loci confirms the strengthening of *cis*-interactions between persistent and transient sites within the clustered enhancer as well as with target *TFF1* and *NRIP1* promoters (Fig 3F and G). These gained or strengthened interactions overlap with the transiently gained ERα peaks within *TFF1* and *NRIP1* LDECs. Overall, these results demonstrate an induced and specific 3D interaction between the transient and the persistent sites, as well as with target gene promoter, in an E2-dependent manner.

**Figure 3.**
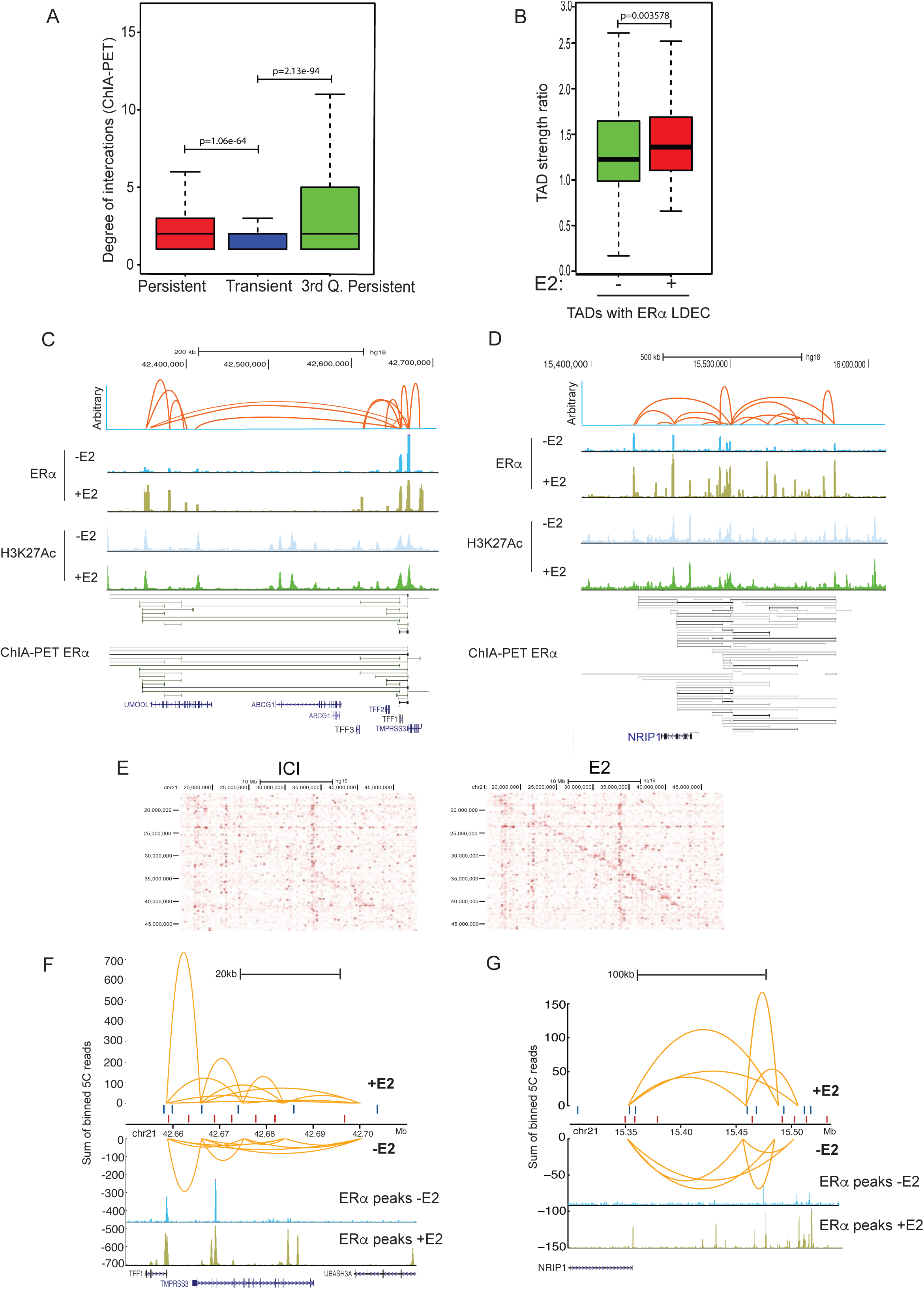
Genomically clustered ERα sites exhibit 3D proximity with each other and with target gene promoter(s) (A) Distributions of the degree of interactions emanating from the persistent, transient and 3^rd^ quantile persistent sites, derived from ERα ChIA-PET data. (B) TAD strength plots from HiC data performed pre- and post-E2 showing the ratio between intra- and inter-TAD interactions emanating from the TADs with LDEC. (C and D) ERα ChIA-PET data plotted for *TFF1* (Left panel) and *NRIP1* (Right panel) LDEC. Orange arches depict the interaction pairs derived from the two replicates, height of loops is arbitrary and do not show the differential strength. ChIA-PET pairs drawn as orange arches are derived from the horizontal lines ranging from light gray (weak interaction) to solid black (strong interaction). (E) 5C matrix from ICI (Left panel) and E2 (Right Panel) conditions showing normalized raw reads. (F and G) Yellow arches on y-axis represent the sum of normalized binned reads as shown for two LDEC, namely, *TFF1* (Left panel) and *NRIP1* (Right panel). Positive y-axis shows the interactions upon E2 treatment whereas negative y-axis corresponds to the interactions in ICI treated cells. The positions of forward and reverse oligos used to derive the plots are shown in red and blue vertical points along the x-axis. The plots are overlaid (bottom) with ERα peaks from-E2 and +E2 ChIP-seq data. All p-values were calculated by either Wilcoxon rank sum test or Wilcoxon signed rank test. The boxplots depict the minimum, first quartile, median, third quartile and maximum without outliers.

### Persistent sites are required for the binding of ERα at neighboring transient sites

Given that the transient sites cluster around persistent sites, we assessed whether persistent sites play a direct role in the emergence of clustered enhancers. We either deleted or blocked persistent site (PS) from LDECs of most inducible genes *TFF1* on Chr21 and *GREB1* on Chr2 using CRISPR-cas9 strategy (Fig S4A, B for *TFF1* PS deletion, S4C and D for *GREB1* PS blocking). ERα occupancy at transient sites around *TFF1* persistent peak was completely lost in ∆PS–tff1 MCF7 cells as compared to the wild-type cells as seen by ChIP-seq of ERα in these cells (Fig 4A, pink highlighted region). Importantly, ERα occupancy at distal sites was unchanged (Fig 4A, Blue highlighted region). The distal sites are in fact the LDEC for *UMODL1* gene. Similarly, we blocked persistent site at GREB1 LDEC using specific gRNA and measured ERα binding at transient sites. Again, not only persistent sites but transient sites also showed loss of ERα binding (Fig 4B and C), supporting essentiality of persistent sites for ERα recruitment at transient sites. We further quantified the effects of enhancer deletion or blocking on the expression of the neighboring genes. The loss of E2-induced expression of *TFF1* was noted (Fig 4D). Interestingly, other genes within the LDEC such as *TFF2 and TFF3* also showed reduced response to E2 signaling in the ∆PS–tff1 cells (Fig 4D); these genes have been shown to physically interact with each other and are regulated by E2 (Quintin et al., 2014, Rafique et al., 2015). Similarly, GREB1 expression was also reduced upon blocking of GREB1 persistent site (Fig 4E). These data strongly suggest the specific and circumscribed effects of persistent site in regulating one (*GREB1* LDEC) or more (*TFF1* LDEC) genes present within LDECs.

**Figure 4.**
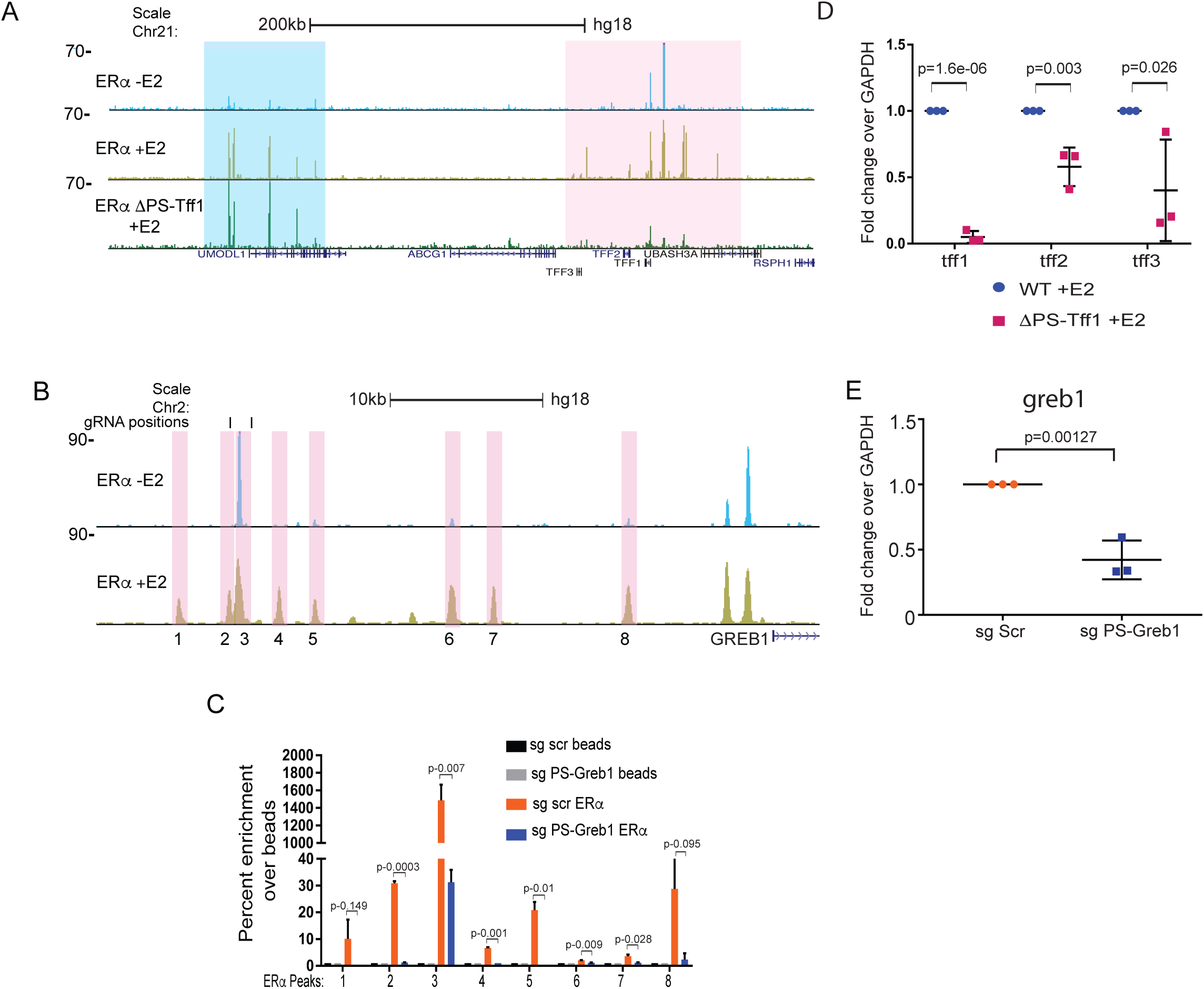
Persistent sites are required for the binding of ERα at neighboring transient sites. (A) Genome browser snapshot of *TFF1* region depicting ERα binding in WT and ∆PS-tff1 lines. First and second ERα ChIP-seq tracks are in untreated and E2 treated WT cells, and the third track represents ERα ChIP-seq in the E2-treated ∆PS-tff1 MCF7 line. Pink and blue highlighted regions represent the *TFF1* and *UMODL1* LDEC respectively. (B) Genome browser snapshot of *GREB1* LDEC exhibiting ERα peaks in untreated and E2 conditions. The vertical highlighted boxes show the regions selected for the measurement of ERα occupancy upon CRISPR blocking of persistent site using gRNAs designed at positions indicated as vertical lines on peak 2/3. (C) ERα occupancy on highlighted regions in panel C upon CRISPR blocking of persistent site shows significant loss of ERα binding on not only the persistent site but also on transient sites. (D) qRTPCRs for TFF1, TFF2, and TFF3 genes in E2 treated WT and ∆PS-tff1 lines (E) RT-PCR exhibiting the loss of GREB1 induction in the cells where *GREB1* persistent site is blocked as compared to the cells transfected with scr gRNAs. QPCR Plots represent data from three biological replicates and each replicate had three technical repeats. Data are plotted as average, SD and p values are indicated. All p-values were calculated by unpaired student’s t-test.

### ERα puncta are formed on LDECs by coalescing

Our results so far point towards ligand-dependent binding of ERα on the EREs near persistent sites and concomitant physical interaction among persistent and transient sites, suggesting spatial crowding of both DNA and ERα in these clusters. We asked if such clusters can be visualized by fluorescence microscopy. Toward this we performed live cell imaging of GFP-ERα upon E2 treatment. Interestingly, the ERα distribution was found to be uniform in untreated cells (Fig 5A) but we observed robust formation of ERα punctate pattern as early as 12-15’ post-E2, formed by coalescing (Fig 5A and 5B, frames from Movie S1 and S2 respectively), number of puncta continued to increase in number and size till 40’ post signaling. Further, to test if the genomic LDECs that emerge in 3D upon ligand stimulation are indeed the dynamic ERα puncta observed in immunofluorescence, we conducted immunoFISH studies on *NRIP1* LDEC as it is one of the densest cluster (Top 10 percentile) in the genome (Fig S5A) and exhibited 3D-proximity within the cluster and target promoters in E2-dependent manner (Fig 3C). BAC clone overlapping LDEC was used. ERα intensity on *NRIP1* loci increased significantly upon ligand stimulation (Fig 5C). Interestingly, ERα puncta intensity at *NRIP1* loci was among the highest intensity ERα puncta that appeared upon E2 stimulation (Fig S5B), likely due to high ERα peak density at this cluster (Fig S5A). These data suggest a model where binding of liganded ERα to transient sites strengthens 3D proximity within LDEC, which can be observed as ERα puncta in the nucleus. To understand the ERα exchange dynamics in these puncta, we performed FRAP experiments on GFP-ERα foci in untreated and cells that were treated with E2 for 1h. Although some puncta were observed even in untreated cells but the recovery of puncta was so robust that a complete bleaching could not be achieved due to rapid exchange of ERα in the puncta (Fig 5D). The recovery of ERα puncta in E2 treated cells was 60%, suggesting a stable ERα puncta formation post-E2 treatment and the loss of fluorescence recovery to this extent has been shown for other punctate patterns (Sabari et al., 2018).

**Figure 5.**
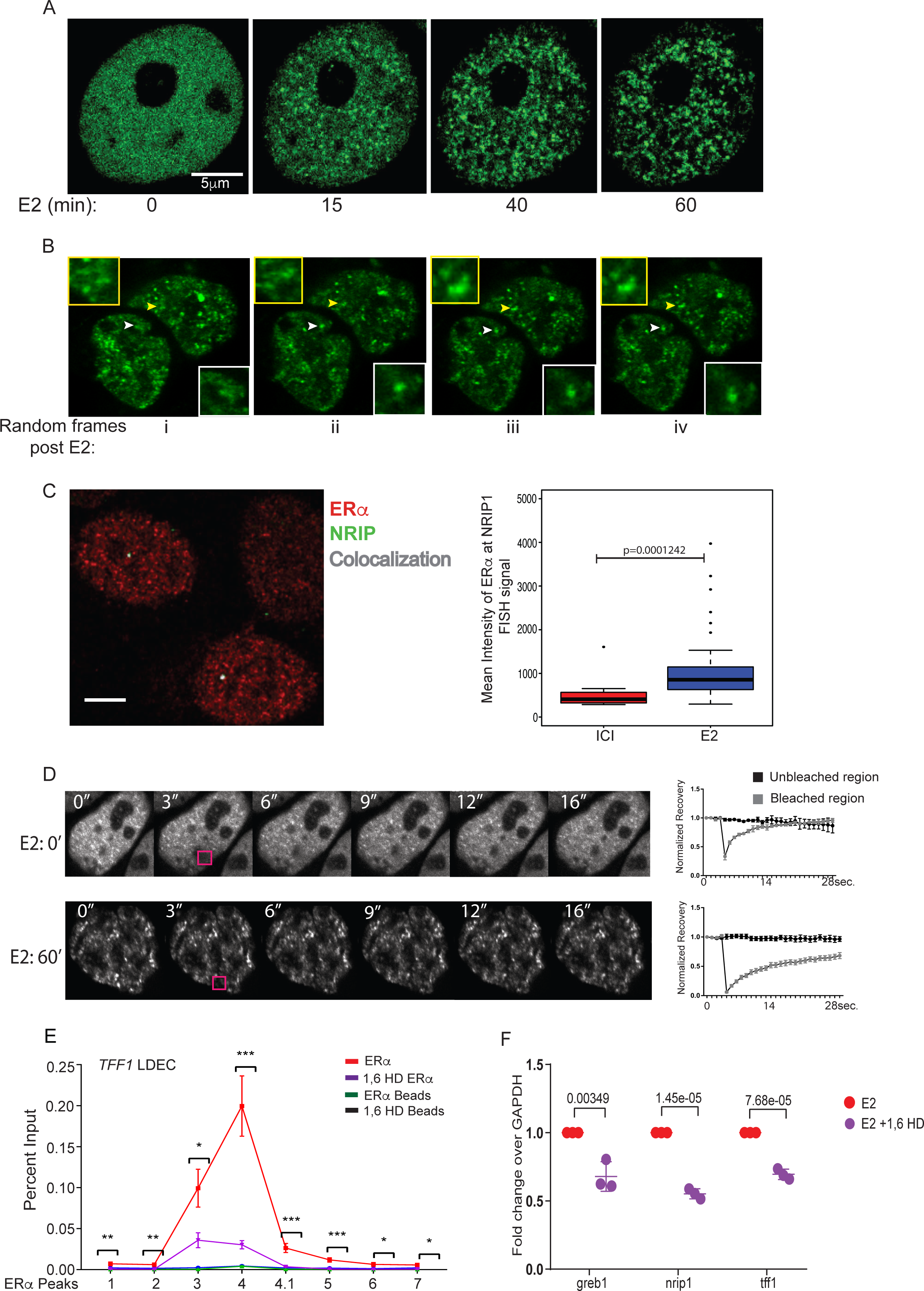
ERα puncta are formed on LDECs by coalescing. (A) GFP-ERα frames from movie showing ERα puncta appear upon E2 signaling during the mentioned time, post-E2 treatment (B) ERα puncta appear upon E2 treatment and puncta of interest are marked by arrows. Insets show the zoomed in images of marked puncta. (C) Immuno-FISH images showing ERα (Green), *NRIP1* FISH signal (Red) and colocalization. Boxplot shows the mean intensities of ERα colocalization in untreated and treated cells. P-value calculated by Wilcoxon rank sum test (D) FRAP recovery of puncta in control cells (upper most panel of images and graph) and at 60’ post-E2 treatment (subsequent panels and graph). Data are as mean +/− SEM. (E) Chromatin Immunoprecipitation shows the levels of ERα at persistent as well as at transient peaks in the *TFF1* LDEC before and after 1,6 HD treatments. (F) E2-dependent expressions of GREB1, NRIP1 and TFF1 is perturbed upon 1,6 HD treatments. QPCR Plots represent data from three biological replicates and each replicate had three technical repeats. Data are plotted as average, SD and p values are indicated. All p-values were calculated by unpaired student’s t-test. The boxplots depict the minimum, first quartile, median, third quartile and maximum.

Recent reports have shown that crowding of protein-occupied regulatory units involve hydrophobic interactions, such as HP1-mediated heterochromatin formation and co-activator mediated condensate formation on enhancers (Larson et al., 2017, Sabari et al., 2018, Boija et al., 2018). To test whether hydrophobic interactions play a role in our observed puncta formation, we treated the E2-pretreated cells with 1,6-hexanediol (1,6 HD) that disturbs hydrophobic interactions. We found that ERα punctate pattern was lost upon 1,6 HD application (data not shown). To recapitulate the loss of puncta at the genomic level, we monitored the occupancy of ERα in cells treated with 1,6 HD. Loss of ERα occupancy was seen both at *TFF1* transient as well as persistent sites (Fig 5E and Fig S5C). The expression of most E2-inducible genes driven by LDECs such as TFF1, GREB1 and NRIP1 was significantly reduced upon administration of 1,6 HD confirming the loss of ERα-clustered enhancers and their functions (Fig 5F). The data suggests that ERα genomic clusters identified indeed correspond to ERα puncta that involve hydrophobic interactions and are formed in a ligand dependent manner.

Med1 and ERα cooperate in gene regulation and were recently shown to cooperate in creating condensates on DNA (Boija et al., 2018). Consistently, we observed E2-dependent binding of Med1 on the clustered enhancers with highest binding on 3^rd^ quantile persistent sites (Fig S6). The data points towards the ERα crowding upon ligand stimulation which corroborates with the 3D genomic interactions that take place within and perhaps between LDECs in E2 dependent manner.

### LDECs exist transiently only during the active phase of signaling

Expression of E2-upregulated genes is known to peak at 45 min to 1 hour before declining at 3 hours post-E2 stimulation (Hah et al., 2011). Further, ERα-bound enhancers have been shown to control most E2-inducible genes (Li et al., 2013, Liu et al., 2014, Hah et al., 2011). We therefore directly assessed whether E2-induced changes in gene expression over the course of signaling are directly driven by E2-induced LDECs, and further, whether the binding dynamics of ERα is different for persistent and transient sites. Towards this, we measured the genome-wide ERα occupancy at various time points post-E2 (5 to 1280 min) using publicly available data (Dzida et al., 2017). We partitioned persistent sites into three quantiles as mentioned before (Fig 1b) to monitor their binding dynamics separately. In general, binding at transient sites is relatively weak (Fig 6A). However, in all categories of sites, a pattern becomes evident, although to different degrees due to varying ERα binding strengths (Fig 6A). We observed an increase in ERα binding strength starting at 5’, peaking at 40’, and then gradually declining to minimal levels at 160’, followed by an unexpected increase in binding strength at later time points. Most evident in the 3^rd^ quantile (strongest) persistent sites, the binding strength recovers almost to the maximum levels ~24 hours post-E2. These results suggest that, LDEC emerge transiently around persistent sites to drive active phase of signaling in terms of target gene expression but they disappear during later phase of signaling which is recapitulated by loss of E2-target gene expression. However, the persistent sites remain ERα-bound as bookmarks (Fig 6B). Given that LDECs correspond to ERα puncta (Fig 5B), we tested if these puncta also follow the similar transient pattern as genomic clusters. We performed ERα immunostaining in cells treated with either ICI for 24h or with E2 for 10’, 60’, 180’, and 24h. Although ERα forms small puncta even in absence of ligand, the overall intensity, size, and the number of puncta increased significantly at 10’, stayed high at 60’ followed by a significant drop at 180’, reaching a minimal level at 24h (Fig 6C). This temporal change in ERα puncta intensity is similar to ERα binding pattern in genome as observed in Fig 6A and B. Further, It is evident collectively from Fig 6A, B and C that even though the persistent sites reappear at 24h (Fig 6A), lack of transient ERα sites around these persistent sites do not allow persistent sites to form the genomic cluster, thereby punctate pattern of ERα disappear dramatically at 3h post stimulation and remains low even at 24h when persistent sites show significant ERα binding (Fig 6A). These data suggest that signaling response is at the peak around 1h however, it drops significantly at 3h post stimulation.

**Figure 6.**
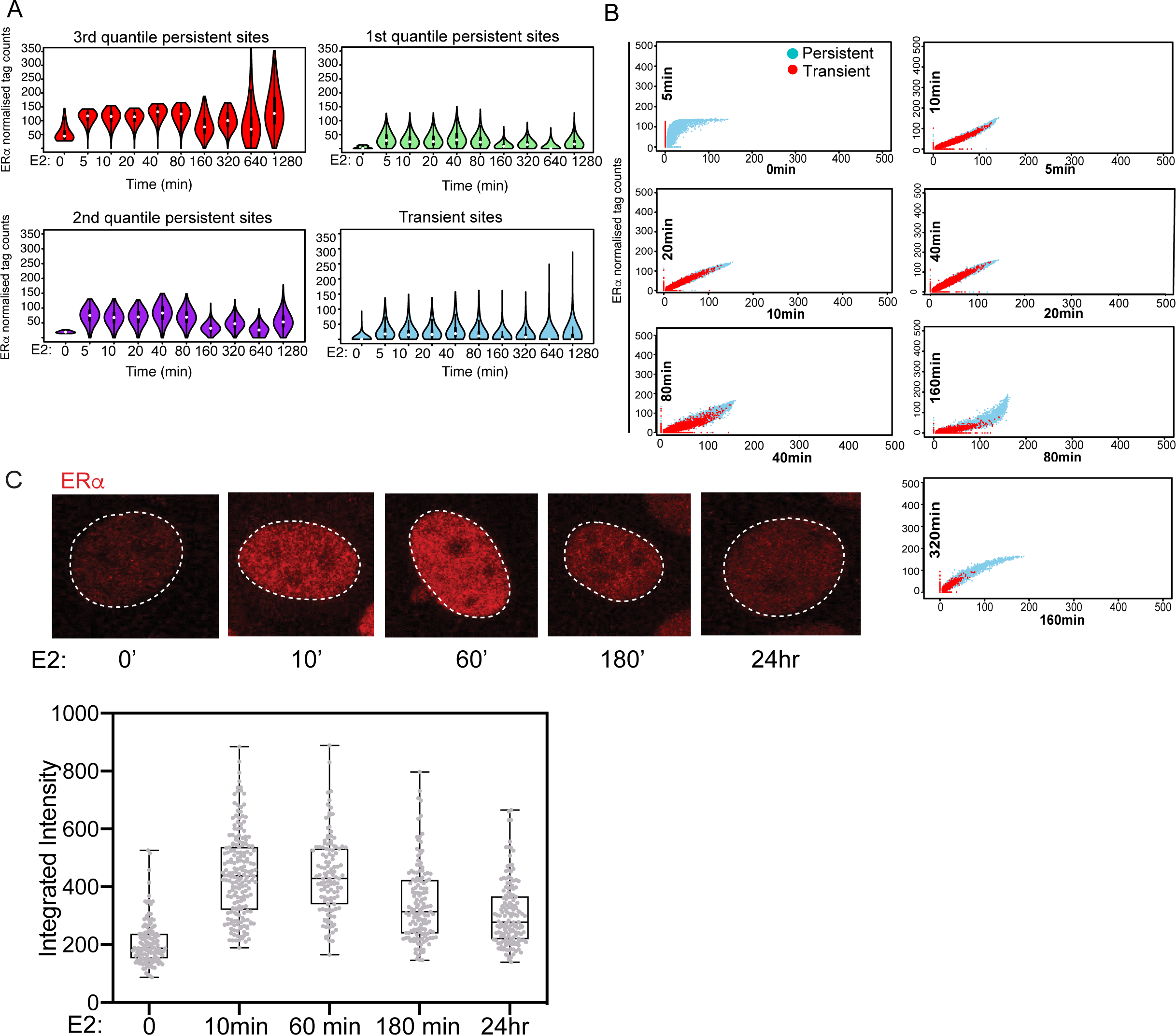
Clustered enhancers exist transiently only during active phase of signaling. (A) ERα binding strength is shown for different categories of ERα peaks at various time points post-E2. The four plots represent 3^rd^ (Top left) 2^nd^ (bottom left) 1^st^ quantile (Bottom left) and transient (Bottom right) ERα peaks. (B) Comparison of binding strengths at consecutive time points for persistent and transient sites reveal an initial surge in binding strength at persistent sites, followed by stably high binding at both persistent and transient sites, followed by loss of transient sites. (C) ERα immunostaining at 0, 10’, 60’, 180’, and 24h post-E2. Bottom panel shows the quantification of ERα integrated intensity at these time points. The boxplots depict the minimum, first quartile, median, third quartile and maximum without outliers.

### Emergence of LDECs is correlated with robust target gene expression

Robust eRNA induction at enhancers is directly linked to robust target gene activation (Li et al., 2013, Hah et al., 2011). Given that ERα clustered enhancers and punctate pattern decline 3h post-E2 treatment, we tested whether expression of eRNA at clustered enhancers undergoes a similar pattern and whether the persistent and transient sites exhibit varying levels of eRNA induction given their differential ERα binding affinity. Toward this, we analyzed time course GRO-seq data (Hah et al., 2011) and found a higher level of eRNAs at strong persistent sites relative to transient and weak persistent sites. Further, the fluctuations in ERα binding at different time points post-E2 stimulation were also reflected in the relative eRNA expression (Fig 7A). Not surprisingly, the persistent sites were the strongest enhancers in the cluster, and the target genes exhibited similar pattern of expression peaking at 40’ and dropping to a minimal level at 180’ post-E2 stimulation (Fig 7B). Conversely, the genes that are away from clustered enhancers but closer to random ERα sites did not show E2-dependent induction. These signaling responses recapitulate the predicted ERα half-life of 3-4 hours in the presence of E2 (Valley et al., 2008, Ried et al., 2003), and the role of ERα degradation kinetics in E2-target gene expression (Nawaz et al., 1999, Lonard et al., 2000, Callige et al., 2005). Similarly, we found that the chromatin-bound fraction of ERα increases from untreated cells to 1h post-E2 treatment but goes down by 3h (Fig 7C), consistent with eRNA, E2-target gene expression, existence of LDECs, and the punctate pattern. This strongly suggests that when ERα protein degrades, the transient sites lose their ERα resulting in loss of clusters reflected by a loss of puncta 3h post-E2. However, ERα binds to persistent sites due to presence of FOXA1 motif at these sites and as expected, FOXA1 knockdown clearly showed the loss of ERα binding at persistent sites (Fig 7D), without affecting the ERα protein levels (Hurtado et al., 2011). Interestingly, FOXA1 knockdown also reduced ERα binding at transient sites lacking FOXA1 motif, supporting a potential crosstalk between the persistent and the transient sites within the cluster (Fig 7D). Our data again suggests that although persistent sites remain bound with ERα, without the ERα binding at the neighboring transient sites they are incapable of driving the target gene expression.

**Figure 7.**
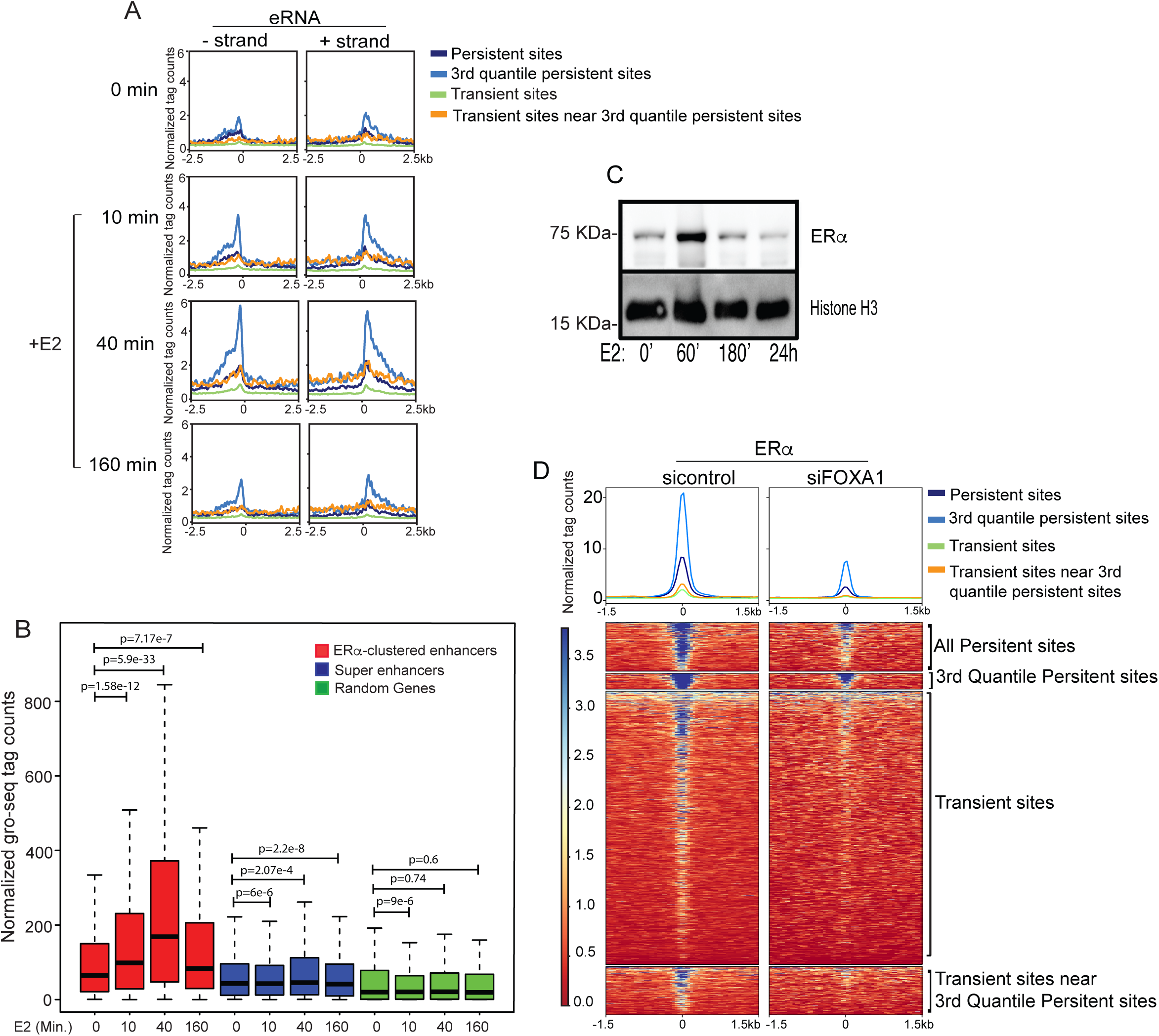
Emergence of ERα clustering is correlated with robust target gene expression. (A) eRNA levels on minus and plus strand show a robust increase in transcription at persistent sites at 40’ post-E2, much more so for the 3^rd^ quantile persistent sites. This is also true to a much smaller extent for transient sites near persistent sites, but not for distal transient sites that are not a part of LDECs. eRNA expression drops to basal level at all sites at 160’, except the strongest persistent sites still show relatively higher levels of eRNAs. (B) Expression of genes closer to all LDEC, H3k27ac super-enhancers and 300 random genes at 0’, 10’, 40’, and 160’ post-E2. (C) Chromatin-bound fractions of ERα at 0, 60’, 180’, and 24hr post-E2 shows the drop in ERα chromatin bound fraction at 180’. (D) Heatmaps showing the binding strength of ERα at various categories of ERα peaks in sicontrol and siFOXA1 transfected cells. All p-values were calculated by either Wilcoxon rank sum test or Wilcoxon signed rank test. The boxplots depict the minimum, first quartile, median, third quartile and maximum without outliers.

**Figure 8.**
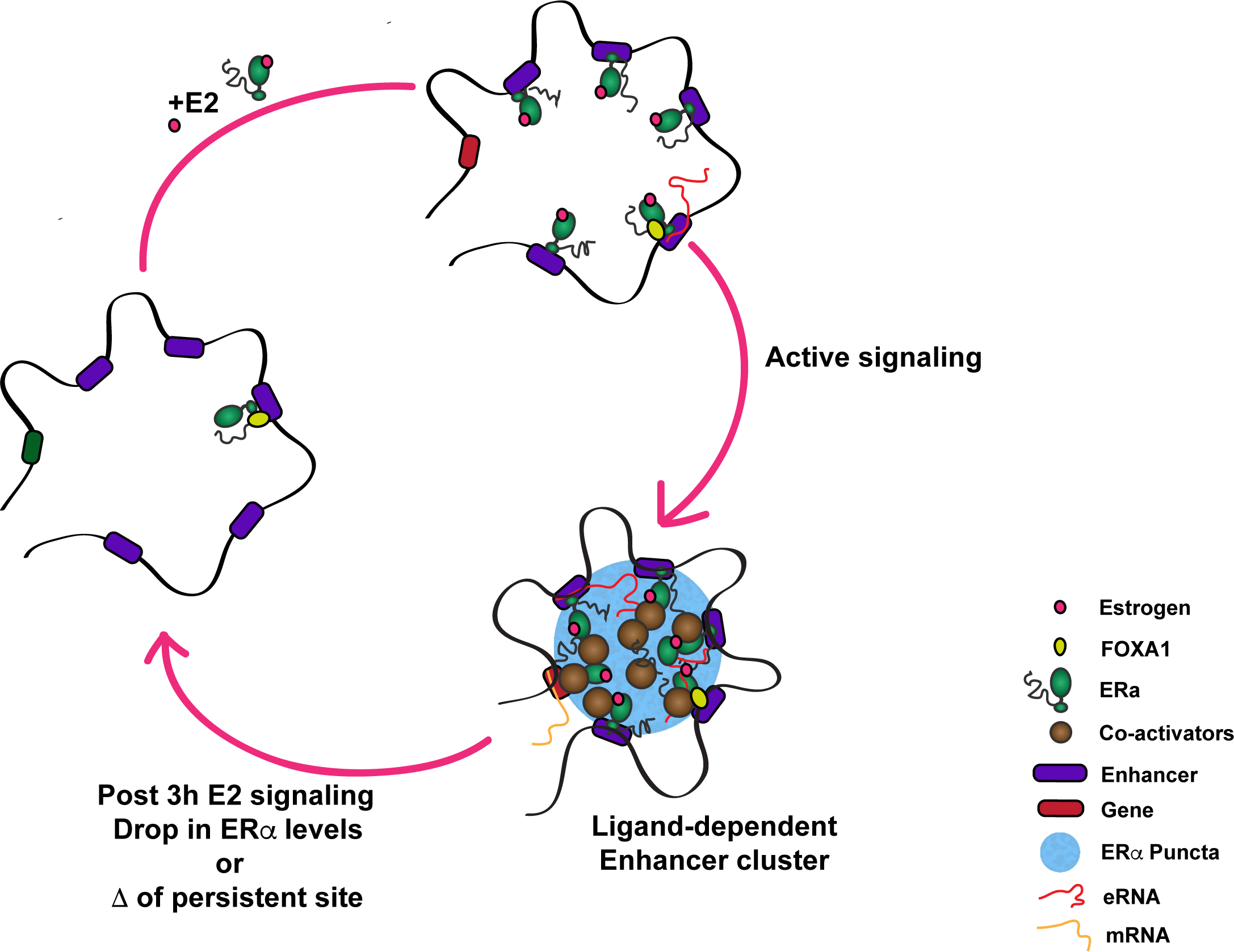
Proposed Model of E2-mediated transcriptional regulation. Model depicts the active and inactive phases of estrogen signaling. During active phase, liganded ERα decorates the EREs closer to premarked-unliganded ERα site. Together, these persistent and transient sites form LDEC in 3D manifesting as ERα puncta, resulting in robust expression of target genes. Upon ERα degradation at 3h or upon deletion of persistent site, these clusters disappear leaving persistent site behind still bound by ERα as bookmark, for next round of ligand stimulation.

## Discussion

### Signaling response is pre-established by an unliganded receptor

Steroid receptors mediate cellular response to hormonal cues by binding to specific enhancers to induce target genes. Our results suggest that, in the case of ERα, a timely and robust response is ensured by pre-marked enhancers bound by unliganded ERα, which nevertheless remain in an inactivated form. However, upon signaling cues, additional liganded ERα binding at EREs in relative proximity to persistent sites is triggered, transiently creating an active cluster of enhancers capable of driving robust gene expression. We found that the presence of FOXA1 motif is a striking feature of pre-marked enhancers (Fig 1D). FOXA1, as a pioneering factor stabilizes the binding of ERα on persistent sites in absence of ligand when levels of nuclear ERα are very low. Upon E2 treatment, when the liganded nuclear ERα levels rise in the nucleus, ERα binds to transient sites albeit with low affinity (Fig 1C). However, at later time points when the level of nuclear liganded-ERα decline by proteolysis again (Fig 7C), FOXA1 still assists binding of ERα at persistent sites but transient sites lose their ERα, restoring the binding pattern of ERα to the pre-treated state.

Despite harboring EREs, unliganded ERα does not bind to transient sites due to low chromatin accessibility (DHS), even when the levels of ERα are raised by ERα overexpression (Fig 5A, 0 min) (Movie 1). However, upon E2 stimulation, liganded ERα alone can penetrate EREs as seen *in vitro* (Gronemeyer, 1991), or it does so by binding with cooperative factors and chromatin remodelers as in the case of glucocorticoid receptors (Grontved et al., 2013, He et al., 2012). However, in absence of persistent site or upon FOXA1 knockdown, transient sites fail to recruit ERα, suggesting a role of FOXA1 in stabilizing the ERα binding not only at persistent sites but also at transient sites within LDECs, likely aided by 3D proximity with persistent sites. Collectively, these results suggest that a framework for signaling is setup by persistent sites via bookmarking the future functional enhancers with FOXA1 and unliganded ERα, which act as a nucleating point for the subsequent ERα binding on the neighboring EREs. Furthermore, chromatin remodeling around persistent sites do not appear to occur as the intermittent H3K27ac sites within *TFF1* LDEC that are not occupied by ERα remain unaffected by E2 signaling (Fig 1E), suggesting that the transient sites within LDECs open up specifically upon E2 signaling. ERα clustered enhancers form a hierarchy where only some of the enhancers control the target gene expression while others seem redundant (Carlton et al., 2017), which is very similar to STAT5-driven super-enhancers (Shin et al., 2016). Our data suggest that perhaps the enhancers that are pre-marked are the functional enhancers.

### LDECs exhibit extensive chromatin looping in ligand-dependent manner

Persistent sites exhibit overall higher degree of interactions (Fig 1A). On the other hand, TADs that contain LDECs exhibit E2-induced increase in intra-TAD interactions relative to their inter-TAD interactions (Fig 3B). This may suggest that in absence of signalling, persistent sites interact with other regulatory sites across different TADs, but upon E2 induction, as transient sites appear bound with liganded ERα, persistent sites redirect their interactions with transient sites favouring more intra-TAD interactions. This E2-dependent shift from long-range to short range interactions is a prominent feature of LDEC. Such E2-dependent alterations in interactions favouring more short-range interactions was also reported recently (Rodriguez et al., 2018, Rafique et al., 2015). However, our study teases apart the specific features of TADs that exhibit such behaviour. These observation also suggest that LDEC formation may not require major changes in chromatin architecture as most alterations are contained within the TADs.

In some instances, LDEC from two neighbouring TADs also exhibit interactions, for example, *TFF1* LDEC shows interaction with *UMODL1* LDEC (Fig 3C), but ERα occupancy at *UMODL1* locus was unaffected by the perturbation of persistent site in *TFF1* LDEC (Fig 4A), suggesting a degree of autonomy and circumscribed influence of each LDEC. Related to this observation, there is a recent interest in understanding whether a portion of chromatin interactions may have functional roles other than transcriptional regulation (Williamson et al., 2014).

### ERα puncta are formed on LDECs by coalescing

While ERα binds to the DNA mostly as a monomer or dimer, as reported recently, it can assume different quaternary structures as well (Presman et al., 2016). This observation is also supported by the fact that dimerization-deficient or DNA-binding mutant of ERα do not form the punctate pattern in the nucleus (Wang et al., 2006). This suggests that interactions among multiple ERα sites within a LDEC could result in ERα-tethered 3D genomic clusters appearing as ERα puncta. Looking closely at Fig 3C, one of the central enhancers in *TFF1* LDEC that is not occupied by ERα does not participate in ERα-bound chromatin network (Fig 3C), suggesting that the intervening DNA that is not bound by ERα may have been looped out allowing the regulation of only tethered DNA in such structures. Along similar lines, intrinsically disordered proteins have been shown to mediate protein-protein interactions while being bound on DNA (Shin et al., 2018). They create genomic clusters by tethering protein bound DNA which results in mechanical exclusion of intervening chromatin fibre. These mechanical principles could define most genomic clusters created in 3D.

We observed Med1 occupancy on LDEC is entirely ligand dependent (Fig 6S) which recapitulates the established interaction between Mediator complex and ligand binding domain of ERα (Kang et al., 2002, Malik and Roeder, 2003). It has been shown that deletion of N-terminal region in ERα exhibits complete loss of punctate pattern in the nucleus upon E2 treatment (Tanida et al., 2015). Further, PONDR analyses suggest that this region of ERα is unstructured. Since, IDRs in transcription factors allow them to phase separate (Liu et al., 2006, Harmon et al., 2017), thus IDR of ERα at its N-terminus and Med1 may allow them to form multivalent homotypic or heterotypic complexes on LDECs. The high density of ERα along with Med1 may allow LDEC to form liquid condensates as shown recently (Boija et al., 2018). Furthermore, persistent sites show robust induction of eRNAs and they interact with Mediator complexes (Lai et al., 2013, Cheng et al., 2018) thus, it is likely that eRNA, Med1, and ERα form RNA protein complexes (RNP) strictly upon E2 stimulation, as shown for other RNA binding proteins (Lin et al., 2015, Banani et al., 2016).

### Loss in signaling response recapitulates the loss of ERα protein

The ERα level is significantly reduced at 3h which is the half-life of ERα in presence of ligand (Fig 7C). The reduced levels of ERα at 3h overlaps with disappearance of LDEC (Fig 6A, B and C) and concomitant loss of eRNA and gene expression (Fig 7A and B) suggesting that upon loss of ERα levels transient sites lose their ERα However, persistent sites regain the binding back due to the presence of FOXA1 at these sites. Interestingly, MG132 pre-treatment of E2-treated cells which stabilizes the ERα levels even at later hours post signaling, allows longer and persistent expression of E2-regulated genes (Fan et al., 2004) conforming that the 26S-proteosome mediated degradation of ERα exerts enhancer-mediated decline in the signaling response.

Together, these observations suggest a mechanism by which unliganded receptor acts as a nucleating point for the new ERα binding in its proximity. Many transient sites along with persistent sites loop with each other to form a 3D complex. These clusters appear to be formed by coalescing of ERα as seen by emerging puncta in E2-dependent manner. Further, the clusters are transient and disappear 3h post signaling coinciding with the loss of eRNA expression and target gene inducibility, clearly depicting the inability of persistent sites in inducing the gene expression on its own unless they form functional units with transient sites. Our work reveals for the first time, enhancer-clustering created by ERα tethering in 3D around unliganded-receptor bound sites, thus forming functional unit that drives active phase of signaling.

## Author contributions

BS, DS, SH, and DN conceptualized the work. DN supervised the work. SH supervised the Informatics work performed by BS. BS, DS, ZI, RJ, RM, SM, UF, AS, KW and SG performed the experiments. Manuscript was written by BS, SH, and DN with input from all authors. Final manuscript was read and approved by all the authors.

## Acknowledgements

DN is a Wellcome-IA Fellow and is also supported by TIFR funding. SH was partly supported by NSF award #1564785 and Fulbright-Nehru scholarship. UF and KW are supported by PhD fellowships from Council for Scientific and Industrial Research (CSIR) India. We thank Sudeshna Majumdar for assisting in ImmunoFISH experiments and Sakshi Gorey for help in CRISPR experiments. Authors would like to thank Dan Larson for his inputs on an earlier draft of the manuscript.

---

**Fig S1.**
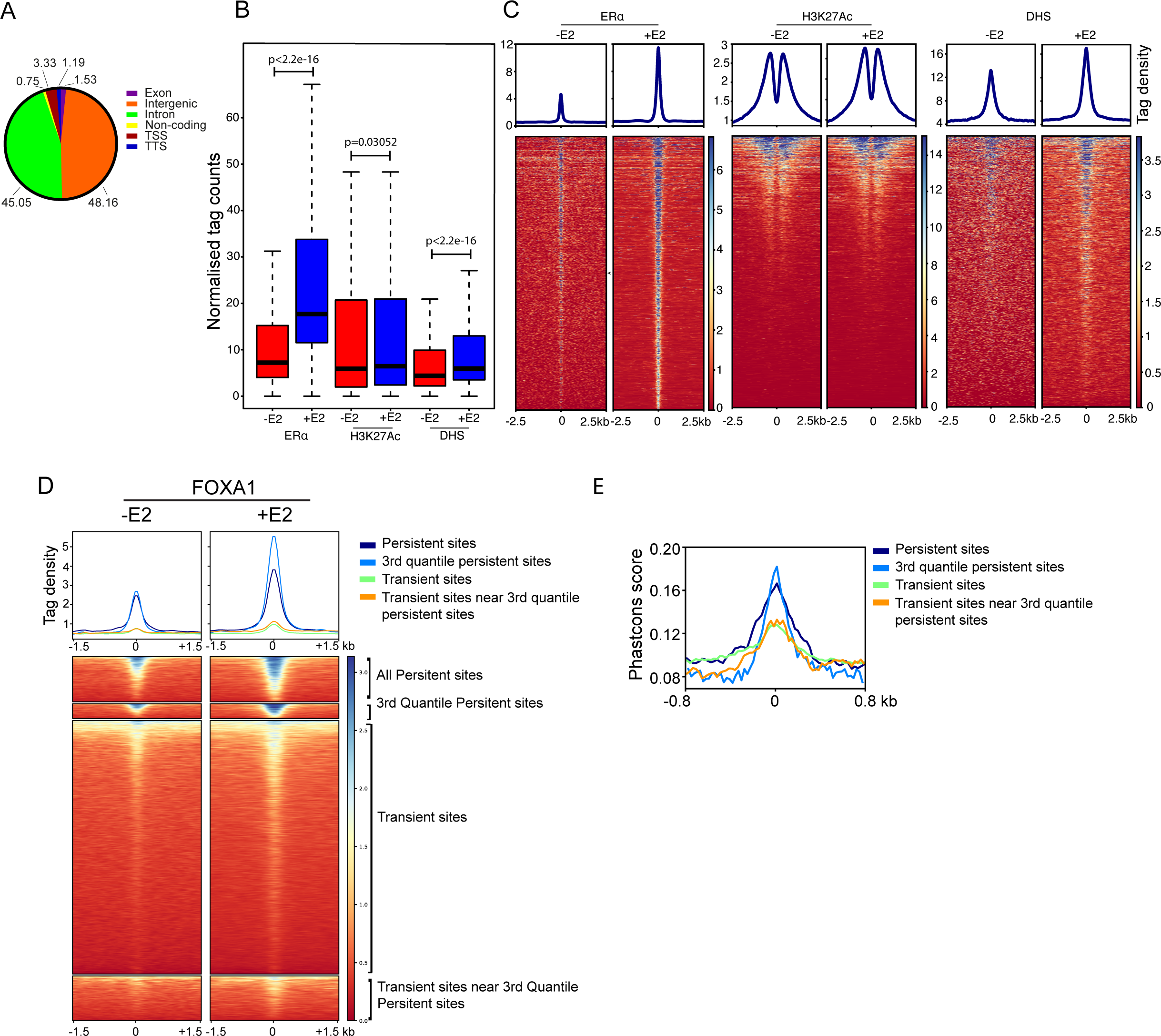
ERα clusters around persistent sites in an E2-dependent manner. (A) Venn diagram shows distribution of ERα on various genomic regions. (B) Strength of ERα occupancy, DHS, and H3K27ac levels at ERα peaks before and after E2 treatments. (C) Heatmaps of ERα, DHS, and H3K27ac 2.5 kb upstream and downstream regions from the center of the ERα peaks in untreated and E2 treated condition. (D) Heatmaps representing the strength of FOXA1 binding in different categories of ERα peaks in treated and untreated cells. Strength was measured at 1.5 kb upstream and downstream of center of ERα peak. (E) Phastcons score of persistent, 3^rd^ quantile persistent, transient, and transient near persistent sites.

**Fig S2.**
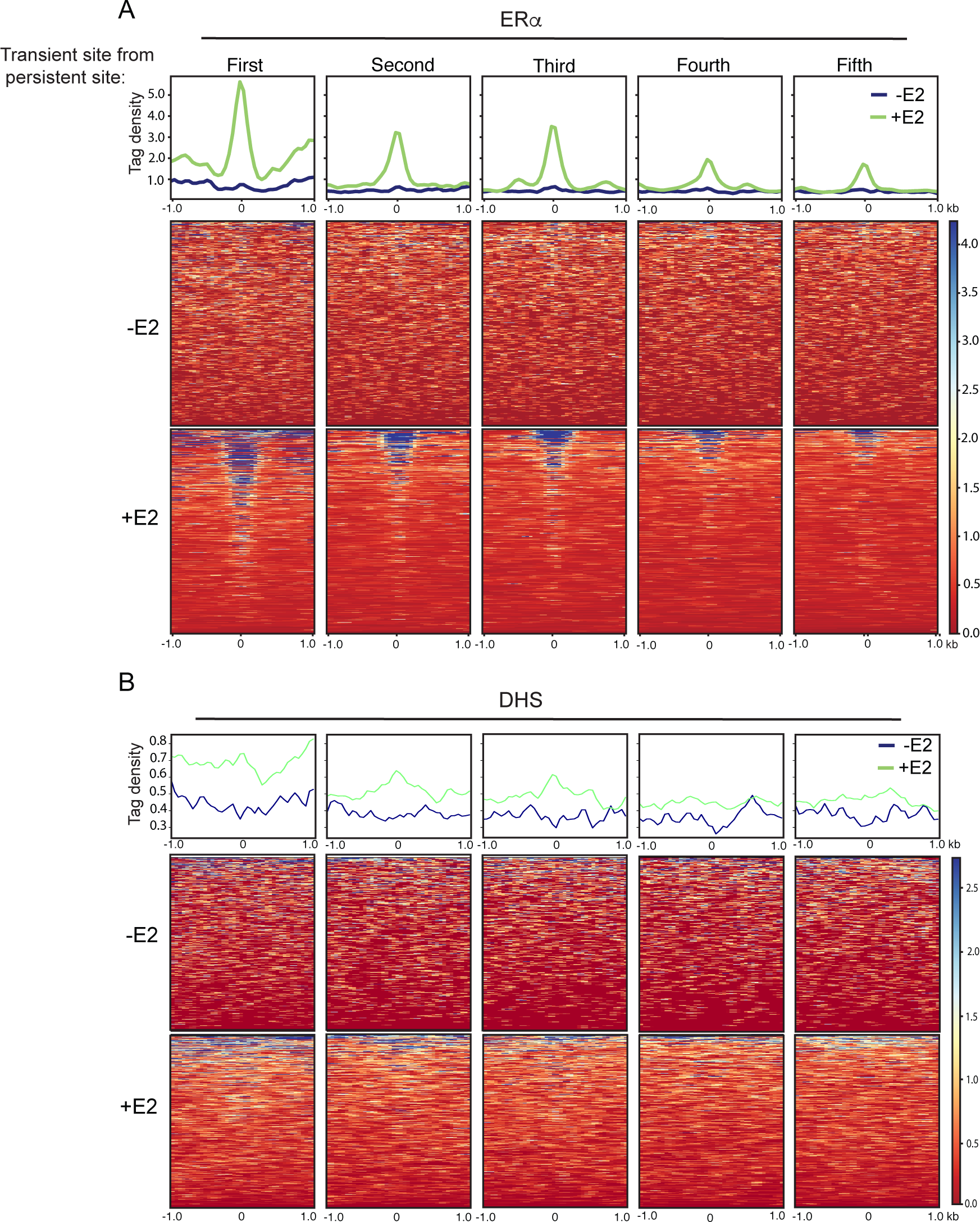
ERα clustered enhancers but not conventional super enhancers control E2 target genes. (A) Heatmaps exhibit the loss of ERα binding strength at every 2 consecutive EREs from persistent site (B) Heatmap shows DHS signal on sites in panel A.

**Fig S3.**
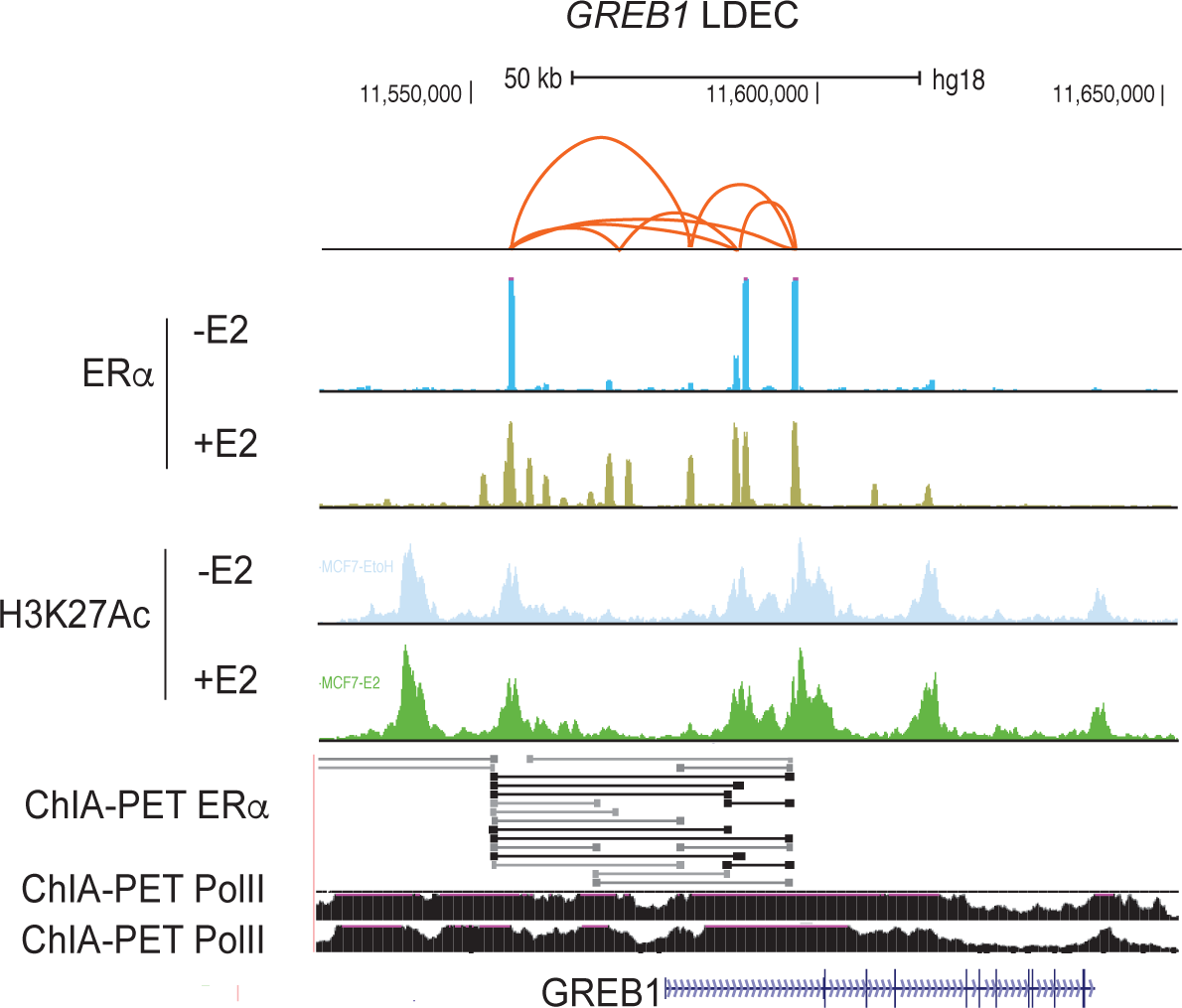
ERα sites within clustered enhancers exhibit 3D-proximity. ChIA-PET data plotted from two ERα ChIA-PET replicates on *GREB1* LDEC as shown in Fig 3C and D.

**Fig S4.**
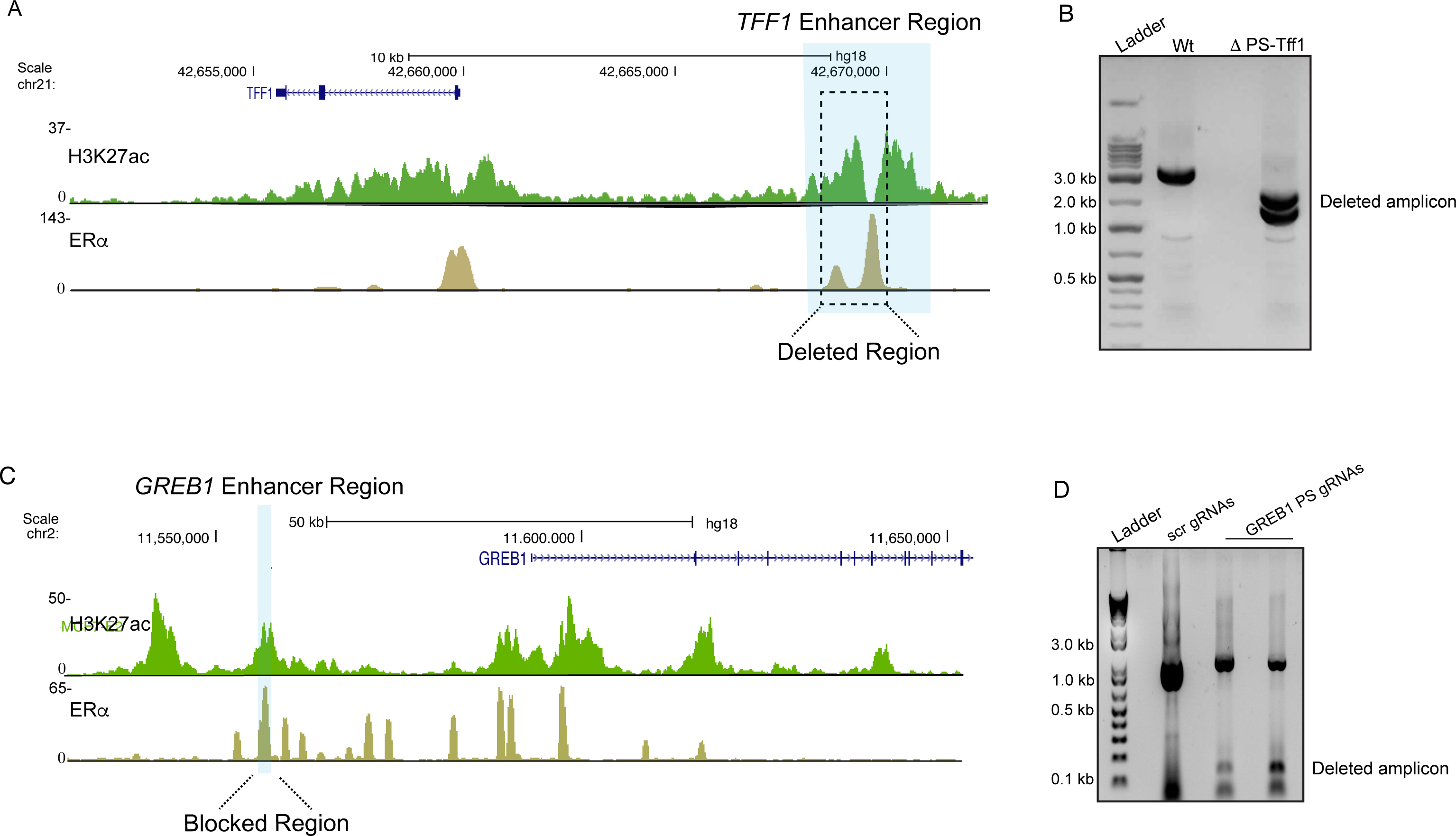
Persistent sites are required for the emergence of transient ERα sites within clustered enhancers. (A) UCSC genome browser snap shot of *TFF1* region showing blue highlighted persistent sites. Dashed line box marks the deleted regions. (B) Surveyor assay using the oligos specific for the region outside the deleted PS. Wt genomic DNA exhibits the larger molecular weight amplicon compared to the amplicon from ∆PS-Tff1 genomic DNA. (C) UCSC genome browser snap shot of *NRIP1* region showing blue highlighted persistent site which was blocked by specific gRNAs. (D) gRNA’s cut the specific region within the enhancer as shown by surveyor assay using oligos outside of blocked region, PCR was performed on population of cells after transfection so larger and smaller both amplicons are seen.

**Fig S5.**
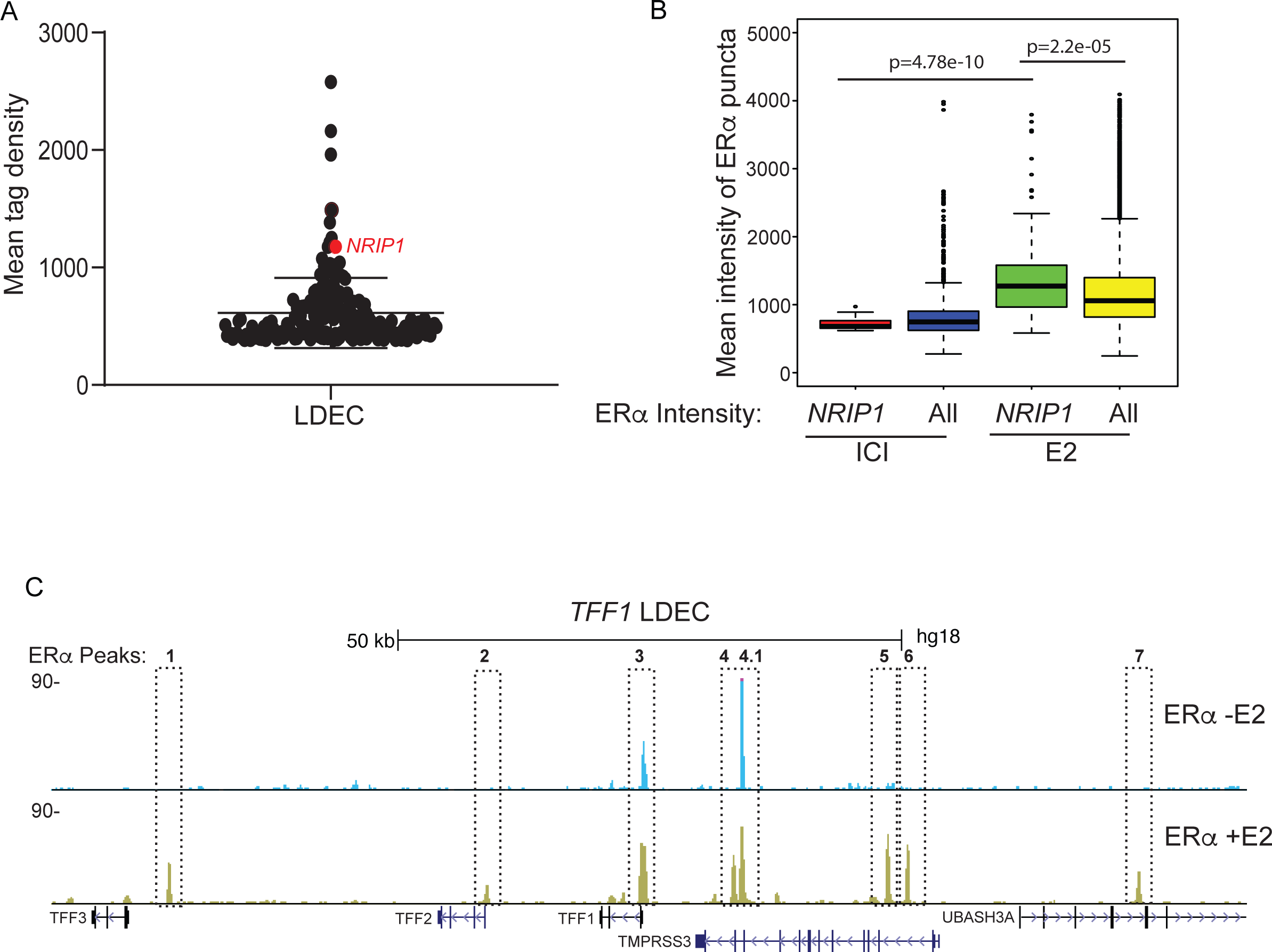
ERα puncta are formed on LDECs by coalescing. (A) UCSC genome browser snapshot of *TFF1* region showing the ERα ChIP-seq peaks in untreated (Top track) and E2 treated (Bottom track) conditions. Dashed boxes mark the regions on which ERα occupancy was measured upon 1,6 HD treatments. (B) Distribution of ERα-bound peak density across all the clusters. (C) Comparison of ERα intensity on all puncta vs. the puncta that overlap with *NRIP1* loci by immunoFISH upon ICI and E2 treatment for 1h.

**Fig S6.**
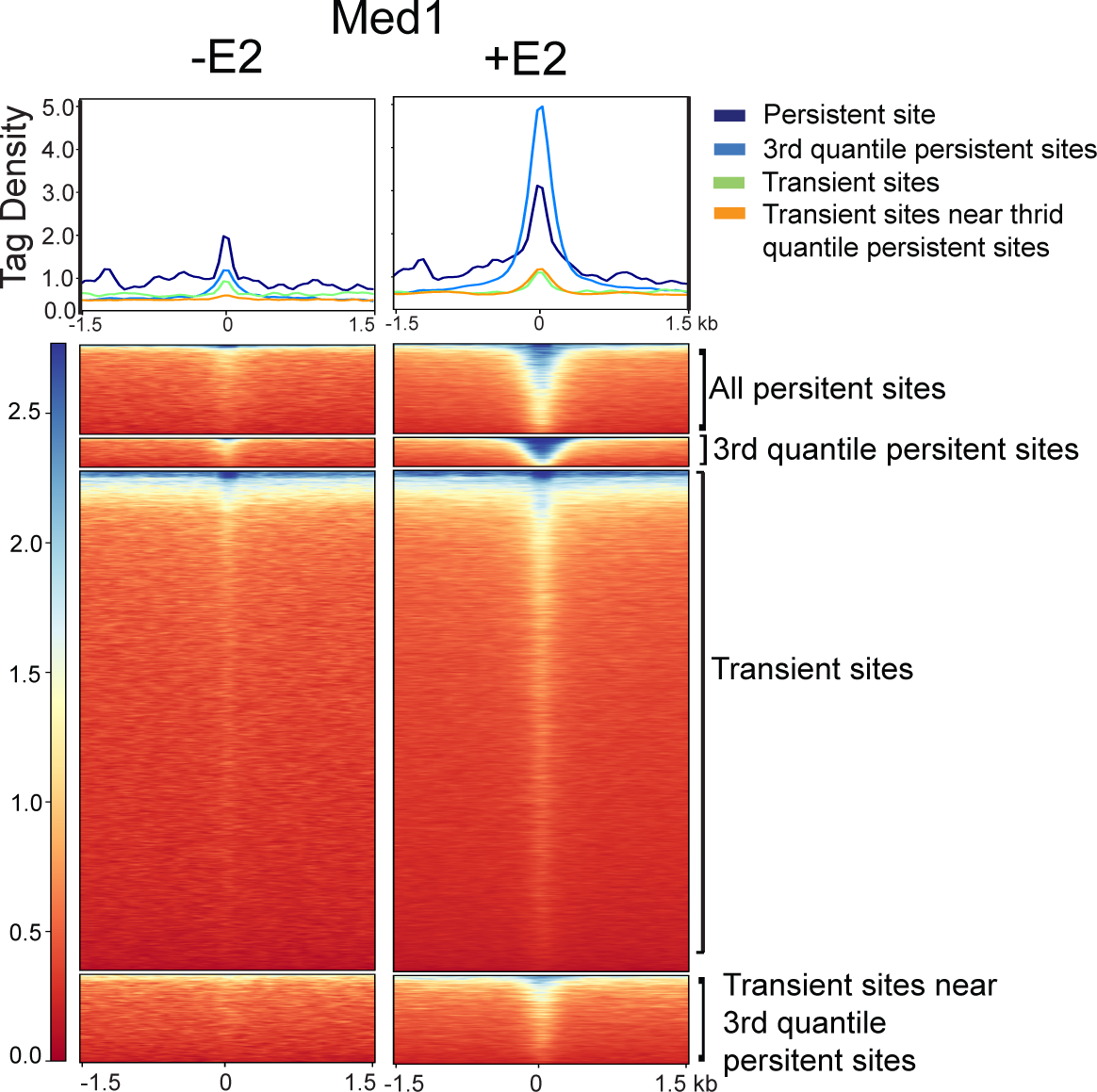
Med1 occupancy is E2 dependent. Heatmaps represent the Med1 binding strength on different categories of ERα peaks in untreated and E2 treated conditions. Tag density was measured on 1.5 Kb upstream and downstream region from center of ERα peaks.

**Movie S1:** Live imaging of GFP-ERα upon E2 stimulation. Rest period 4.8sec, exposure 3.2 sec. total interval 8 sec. Three movies taken for different periods were stitched together.

**Movie S2:** Live imaging of GFP-ERα post E2 stimulation. Rest period 4.8sec, exposure 3.2 sec. total interval 8 sec.

## Materials and Methods

### Cell Culture

MCF7 cells were obtained from ATCC. They were cultured in High glucose DMEM media (Invitrogen) at 37°C and 5% CO_2_ conditions in humidified chamber. For ligand stimulation, cells were grown in stripping media containing high glucose DMEM without phenol red and 10% charcoal stripped FBS for three days. On the third day cells were treated with β-estradiol (E2758, Sigma-Aldrich) at 10nM concentration for various periods as mentioned in the respective figures. For untreated control, cells were either treated with equal microliters of ethanol on the third day or with ERα inhibitor ICI182780 (1047, Tocris Biosciences) at 10nM concentration for 24 h after two days of stripping.

### 1,6 Hexanediol treatments

Cells pretreated with either E2 or vehicle for 30 min were then treated with 5% 1,6 HD for 30 min in the same E2/vehicle containing media. Cells were then cross-linked to be processed for ChIP or were directly subjected to RNA isolation using Trizol method (Invitrogen) and subsequently cDNA synthesis was performed.

### Chromatin Immunoprecipitation and qPCRs

ChIP protocol was followed as described in (Li et al., 2013). Briefly, 5 million MCF-7 cells were cross-linked using 1% formaldehyde for 10 min at room temperature. Formaldehyde was quenched by 125µM Glycine for 5 min at RT with gentle rotation. Cells were scrapped in cold PBS, pelleted at 2000 rpm for 5 min at 4^°^C. Pellet was dissolved in lysis buffer (50µM Tris HCl pH 7.4,1%SDS, 10µM EDTA and protease inhibitors), sonicated using Bioruptor Pico (Diagenode) to obtain fragments of 500 bp. Sonicated lysate was spun to remove the insoluble debris. Supernatant was diluted 2.5 times with dilution buffer (20mM Tris HCl pH7.4, 100µM NaCl, 2µM EDTA, 0.5% TritonX100 and protease inhibitors). 100 µg chromatin was taken for each IP, 1 µg ERα (sc-8002, Sigma-Aldrich and ab32063, Abcam), antibody was added to bind with complexes overnight at 4^°^C with gentle rocking motion. 15 µl 50% slurry of Protein G dynabeads (10004D, Invitrogen) were added for 4 hours and complexes were captured on magnetic rack, and the complexes were washed three times. The eluted complexes were reverse cross-linked overnight at 65^°^C followed by purification with phenol:chloroform:isoamylalcohol. Purified chromatin was dissolved in 100 µl of TE (pH.8) buffer and 4 µl of chromatin was used per q-PCR reaction on CFX96 touch real time PCR (Bio-rad Laboratories). q-PCRs were carried out in three technical replicates and at least three biological replicates were performed. The fold changes were calculated by 2^-∆∆ct^ method or percent over input was calculated as described in https://www.thermofisher.com/in/en/home/life-science/epigenetics-noncoding-rna-research/chromatin-remodeling/chromatin-immunoprecipitation-chip/chip-analysis.html.

### ChIP-seq Library preparation and data analysis

Library preparation was performed as per manufacturer instruction for NEB Next ChIP-seq library preparation Kit (New England Biolabs). Briefly, at least 10 ng chromatin was subjected to end repair, T tailing and adapters compatible with illumina sequencing platform were ligated. The amplified library was size-selected using AMPure XP beads to enrich fragments from 300 to 500 bp in size. 12 pico moles of library was used for cluster generation and sequencing was performed in 1×50 bp format on Hiseq 2500 (illumina Inc.).

The sequenced reads were aligned to hg18 assembly using default Bowtie2 options. Tag directories were made from the aligned reads to identify ChIP-seq peaks using HOMER. A 200bp sliding window was used to identify narrow peaks which are characteristic of transcription factor peaks. The common artifacts from clonal amplification were neglected as only one tag from each unique genomic position was considered. The threshold was set at a false discovery rate (FDR) of 0.001 determined by peak finding using randomized tag positions in a genome with an effective size of 2 × 10^9 bp. For ChIP-seq of histone marks, seed regions were initially found using a peak size of 500bp (FDR<0.001) to identify enriched loci. Enriched regions separated by <1kb were merged and considered as blocks of variable lengths. The read densities as bed graph files were calculated across the genome and this track was uploaded to UCSC genome browser.

### ERα-Clustered enhancers

ChIP-seq peaks called (described above) were selected if they were within 20kb distance of each other. Clustered enhancers were considered as clusters of three or more such peaks.

### Super-enhancer Calling

Super-enhancers were identified using the ROSE (Rank Ordering of Super-enhancers) algorithm (https://bitbucket.org/young_computation/) using the aligned ChIP-seq reads as input with parameters-s 15000.

### Phast-Cons Analysis

PhastCons scores (scores for multiple alignments of 99 vertebrate genomes to the human genome) across the human genome as a bigwig file was obtained from UCSC genome browser. This was then used to plot PhastCons scores across different regions.

### Gro-seq analysis

The reads were aligned to hg18 assembly using default Bowtie2 options. The reads were counted from the region between +1kb of gene promoter proximal end to 13kb of the gene body using HOMER for each gene. The read densities were calculated similar to that of the ChIP-seq analysis in a strand specific format.

### 5C library preparation and data analysis

5C oligos were designed in alternate orientation (Sanyal et al., 2011) on ERα peaks overlapping with very strong enhancers on chr21 as identified in (Liu et al., 2014), sequences of 5C oligos are mentioned in Table S1. 5C was performed in duplicates using BamHI restriction endonuclease as reported earlier (1 and 2). 5C reactions were performed in ICI and 1h E2 treated cells as reported (van Berkum et al., 2009 and Ferraiuolo et al., 2012). Number of reads for each 5C in ICI are shown in the Table S2 and the Pearson correlation was more than 0.95 for ICI as well as E2 libraries. Raw sequencing reads were mapped to an artificial genome of all possible combinations of forward and reverse primers using Bowtie2. The uniquely mapped reads were then used to build a matrix of interactions between all the reverse and forward primers. The intra-chromosome 21 primers interactions were then normalized to their interactions with chr16 gene dessert for each interaction combination. The normalized read counts for the forward and the reverse oligos falling in a given region were binned and sum of the reads was plotted as arches using Sushi (http://www.bioconductor.org/) in Fig 3g and h. Sequences of oligos used in the Fig 3g and h are mentioned in Table S1.

### ChIA-PET data analysis

ERα ChIA-PET in MCF-7 interaction files were obtained from ENCODE. All three replicates were merged to obtain common ends in all replicates using pairtopair tool from BED tools. Interaction ends were annotated as the different classes of ERα sites using pairtobed BEDtool. The number of interacting partners from each annotated interaction end was defined as the degree of the ERα site and the degree was then plotted for the different classes.

### Immunostaining

Immunostaining was performed as mentioned in (Notani et al., 2010). Briefly, MCF-7 cells were grown on TC coated coverslips in phenol free media containing charcoal stripped serum for 24 hrs. Cell were treated with 10nM E2 or ethanol as vehicle for various time points before fixing them with 4% paraformaldehyde or methanol for 10 minutes followed by permeabilization for 10 min with 0.1% Triton-X-100 in PBS followed by blocking with 1% BSA for 15 min. Coverslips were incubated with ERα antibody (sc-8002, Sigma-Aldrich Corporation) followed by incubation with secondary antibodies conjugated with fluorophores (Life sciences Inc.). Coverslips were treated with 1nM DAPI for 2 min and mounted using vectashield antifade mounting media (H-1200, Vector Laboratories).

### Cell Imaging

Cells were imaged using PLAPON 60x/1.42 oil objective of Olympus FV3000 microscope. Images were taken with intervals of 1.1 seconds.

### Immunofluorescence (IF) coupled with DNA-FISH and the analysis

IF/DNA-FISH was performed following the protocol as described in (Gayen et al., 2015). Briefly, Cells were permeabilized through treatment with cytoskeletal extraction buffer (CSK:100 mM NaCl, 300 mM sucrose, 3 Mm MgCl2, and 10 mM PIPES buffer, pH 6. 8) containing 0.4% Triton X-100 (SRL, #64518) and fixed with 4% paraformaldehyde. For IF, cells were washed 3X in PBS for 3 min each and then incubated in blocking buffer (0.5 mg/mL BSA, in 1X PBS with 0.2% Tween-20) for 30 min at 37°C in a humid chamber. Following blocking, cells were next incubated with ERα antibody (sc-8002, Sigma-Aldrich Corporation) for 1 hr. The samples were then washed 3X in PBS/0.2% Tween 20 for 3 min. and incubated with fluorescently-conjugated secondary antibody (Alexa Fluor, Invitrogen) for 30 min. After three washes in PBS/0.2% Tween-20 for 3 min each, cells were processed for DNA FISH. For DNA FISH, cells were refixed with 1% paraformaldehyde containing 0.5% Tergitol and 0.5% Triton X-100. Next, dehydration was done through wash with ethanol series (70%, 85%, and 100% ethanol, 2 min each) and air dried for 15 mins. The cells were then treated with RNase A (1.25 ug/µl) at 37^°^C for 30 min. The cells were again dehydrated and air dried as described above. The samples were then denatured in a prewarmed solution of 70% formamide in 2X SSC on a glass slide stationed on top of a heat block set at 95^°^C for 11 min followed immediately by dehydration through a −20°C chilled ethanol series (70%, 85%, 95%, and 100% ethanol, 2 min each). The cells were then air dried and hybridized with probe for overnight at 37^°^C. The next day, the samples were washed twice with prewarmed 50% formamide/2X SSC solution at 39^°^C and 2X with 2X SSC, 7 min each. The dsDNA FISH probes were made by randomly-priming DNA templates using BioPrime DNA Labeling System (18094011, Invitrogen). Probes were labelled with Fluorescein-12-dUTP (Invitrogen) Cy3-dUTP (ENZO life science).

Following BAC probe (BACPAC Resources) was used:

NRIP1: RP11-213G23 (hg18, chr21:15245033-15430733)

### ImmunoFISH intensity analysis

The Immunofluorescence coupled DNA-FISH slides were imaged using PLAPON 60x/1.42 oil objective of Olympus FV3000 microscope with the same settings. IMARIS was used for the colocalisation analysis. ERα immunofluorescence and NRIP1 FISH signal was used to identify spherical spots of 0.8um and 1um diameter respectively. Spots were identified using the same intensity thresholds. Spots colocalising with each other were identified with distance threshold of 1 pixel and classified as colocalised and non-colocalised spots. The intensity of ERα signal at the ERα and NRIP1 spots was calculated and plotted.

### FRAP quantification

GFP-ERα (Addgene #Plasmid 28230) was overexpressed in MCF7 cells in phenol free media with charcoal stripped serum. Cells were treated with Ethanol (Vehicle) or with E2 (10 nM) for one hour post 24h of transfections. For 1,6 HD treatments, 5% 1,6 HD was given to transfected cells that were pretreated with E2 for 30 min to continue other 30 min but along with 1,6 HD. FRAP was performed on Olympus FV3000 microscope with 488nm laser. Bleaching was performed over an area of 1!" using 100% laser power and images were collected every two seconds. The intensity of the photo-bleached ROI was calculated across all the frames in Fiji (Schindelin et al., 2012). Background intensity was subtracted from the value of intensity at the ROI at each frame. This value was then normalized to the whole cell intensity and plotted. The same value is calculated for an unbleached ROI in the same way as well and plotted to control for bleaching due to image acquisition (Sprague et al., 2004).

### RNA Isolation and cDNA synthesis

RNA was isolated using Trizol (Invitrogen) as per the manufacturer protocol. 1 µg of RNA was taken for cDNA synthesis using random hexamer as per manufacturer guideline (Superscript IV RTPCR kit, Life Technologies). cDNA was subjected to real time PCR in triplicates in CFX96 touch real time PCR (Bio-rad laboratories) using oligos mentioned in Table S3. Fold changes were calculated by 2^-∆∆ct^ formula. GAPDH or b-actin expression was taken as control for fold changes in gene expression.

### HiC-Analysis

The raw reads were mapped to hg18 assembly using bowtie 2. The HOMER program makeTagDirectory was first used to create tag directories with tbp 1. Data was further processed by HOMER in order to remove small fragment and self-ligations using makeTagDirectory with the following options: -removePEbg-removeSpikes 10000 5. Next, findTADsAndLoops.pl was used to obtain overlapping TADs, produced at a 20kb resolution with 40kb windows. Smallest overlapping TADs were selected and intersected with ERα clusters. Strength of these TADs were obtained as Inclusion Ratios (the ratio of intra-TAD interactions relative to interactions from the TAD to the surrounding region (both upstream and downstream of the TAD to regions the same size as the TAD) across the different conditions using findTADsAndLoops.pl score option.

### Nuclear lysate fractionation and Immunoblotting

Cells were washed twice in phosphate buffered saline (1XPBS) and cell pellet was carefully suspended in 200ul of SFI Buffer (100mM NaCl, 300mM Sucrose, 3mM MgCl2, 10mM PIPES [pH 6.8], 1mM EGTA, 0.2% Triton X-100,) with PIC. Cells were incubated for 30 mins at 4^°^C and then pelleted down at 2900 rpm for 5 min at 4^°^C, and the supernatant was collected (Soluble fraction). The pellet was carefully washed twice in SFI buffer and chromatin bound fraction was extracted by adding 2X protein loading dye to the pellet. Fractions were loaded on 15% SDS PAGE and western was done for ERα (sc-8002 Santa Cruz Biotechnology) GAPDH (sc-32233 Santa Cruz Biotechnology), Histone H3 (H0164, Sigma-Alderich). Cellular lysates prepared were separated on 15% SDS-PAGE gels. The protein transfer was carried out in Tris-glycine buffer at 30V for 1 hour on ice using 0.45µ PVDF membrane (Millipore). Membrane was blocked in 5% non-fat dry milk made in TBST and further incubated for three hours with ER alpha (sc-8002 Santa Cruz Biotechnology) GAPDH (sc-32233 Santa Cruz Biotechnology), Histone H3 (H0164, Sigma-Alderich) antibodies followed by stringent washings. Membranes were probed with HRP-conjugated secondary antibody (Bio-rad Laboratories). Signal amplification was performed using ECL substrate (RPN2106, GE Healthcare). Images were captured on Image Quant LAS4000 with CCD camera.

### gRNA design and cloning

We used gRNA design tool at crispr.mit.edu to design the gRNAs for ERα peak in the GREB1 and TFF1 persistent and transient sites (Table S3). The gRNAs were cloned in a customized PX459 vector (pSpCas9(BB)-2A-Puro V2.0, Addgene # 62988) (Ran et al., 2013) was a gift from Feng Zhang. The Cas9 enzyme cassette was removed from PX459 by digesting it with Xba1 and Not1 followed by blunt end ligation. The cloning of the gRNAs in the customized PX459 plasmid was performed as per the Zhang Lab general cloning protocol.

### Deletion of persistent sites

Cells were transfected with cas9 plasmid and gRNAs constructs described above. Transfected cells were kept under puromycin selection for 48h post 24h of transfection. After two days of selection, cells were diluted at the density of 0.5 cell/100 µl and 100 µl was dispensed into each well of a 96 plate. Wells containing single cells were identified under microscope and marked. Media was changed every 5 days till the colony appeared. Colonies from single cells were screened for the homozygous deletion. Second round of gRNA transfection was done on heterozygous line that was obtained from the first round of cas9 experiment.

**NGS data sets used in this study are mentioned in Table S4.**

---

**Table S1:**
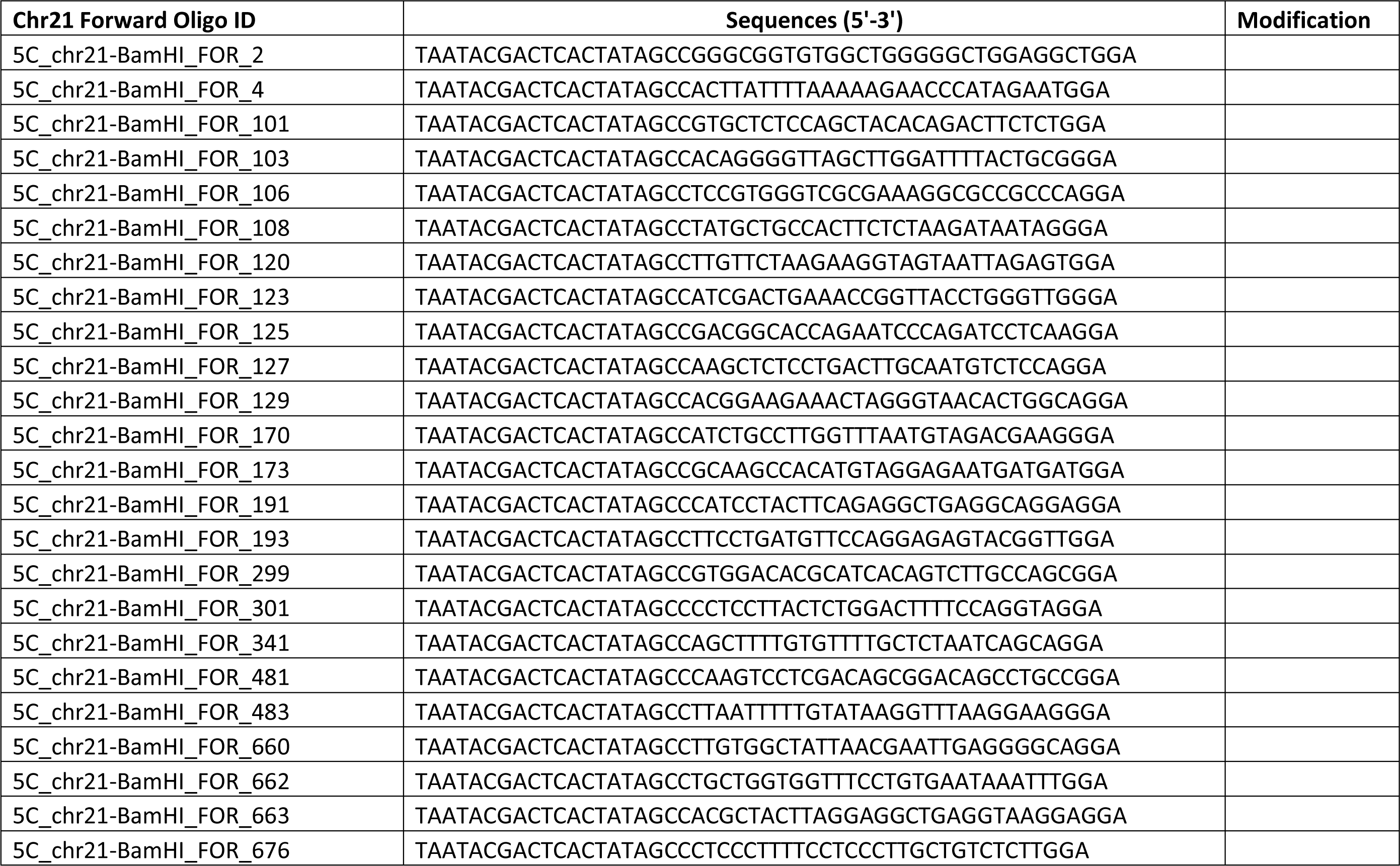
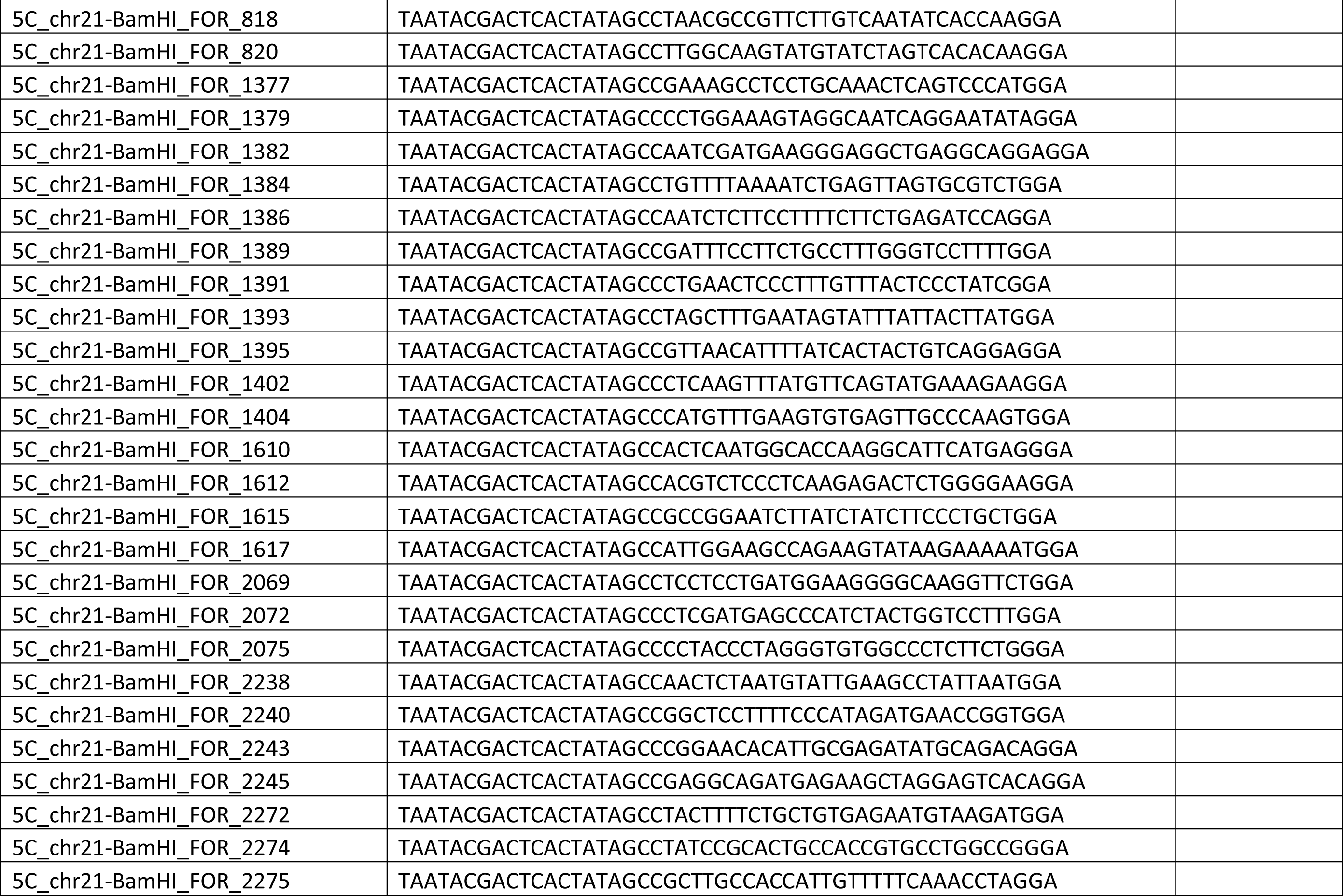
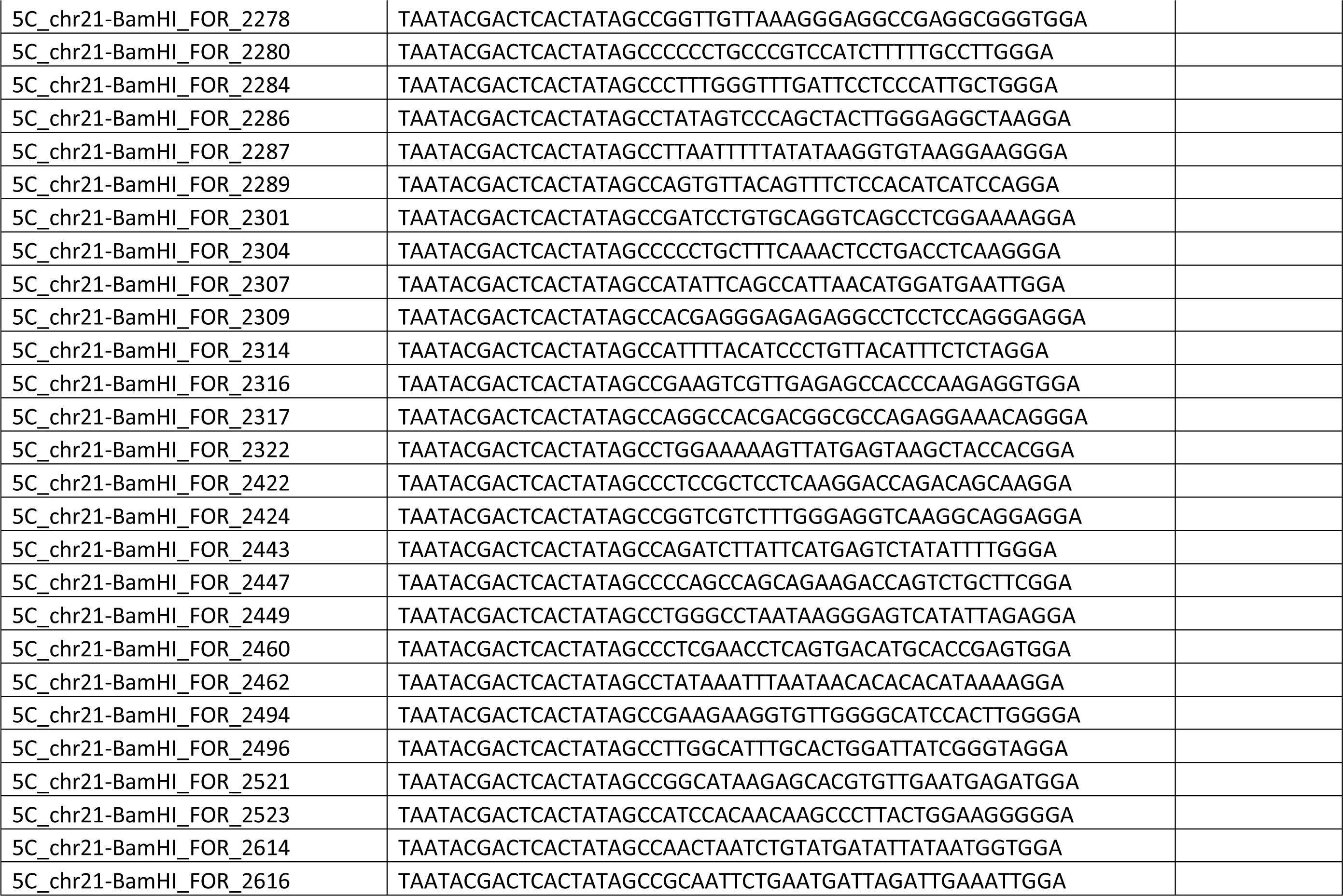
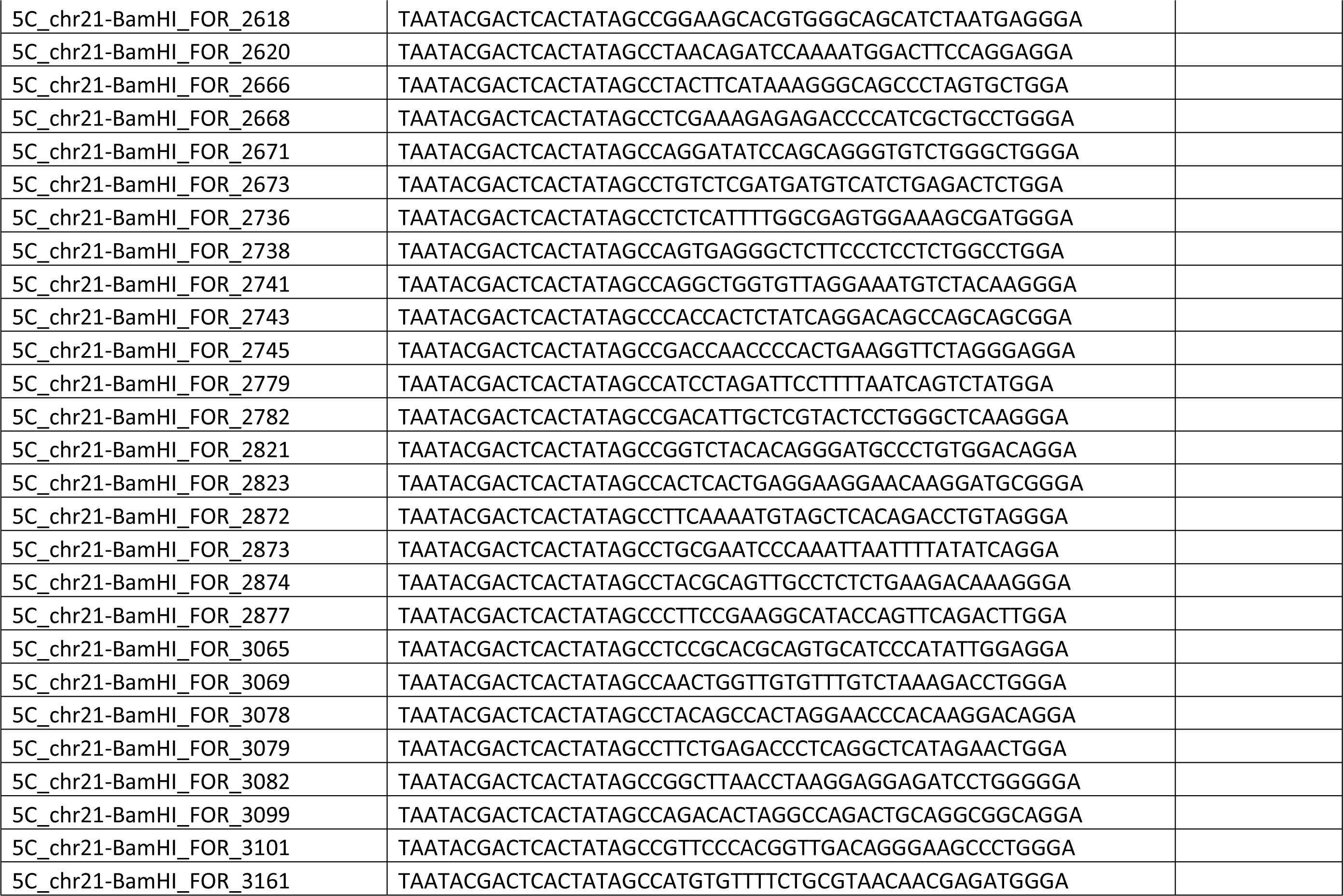
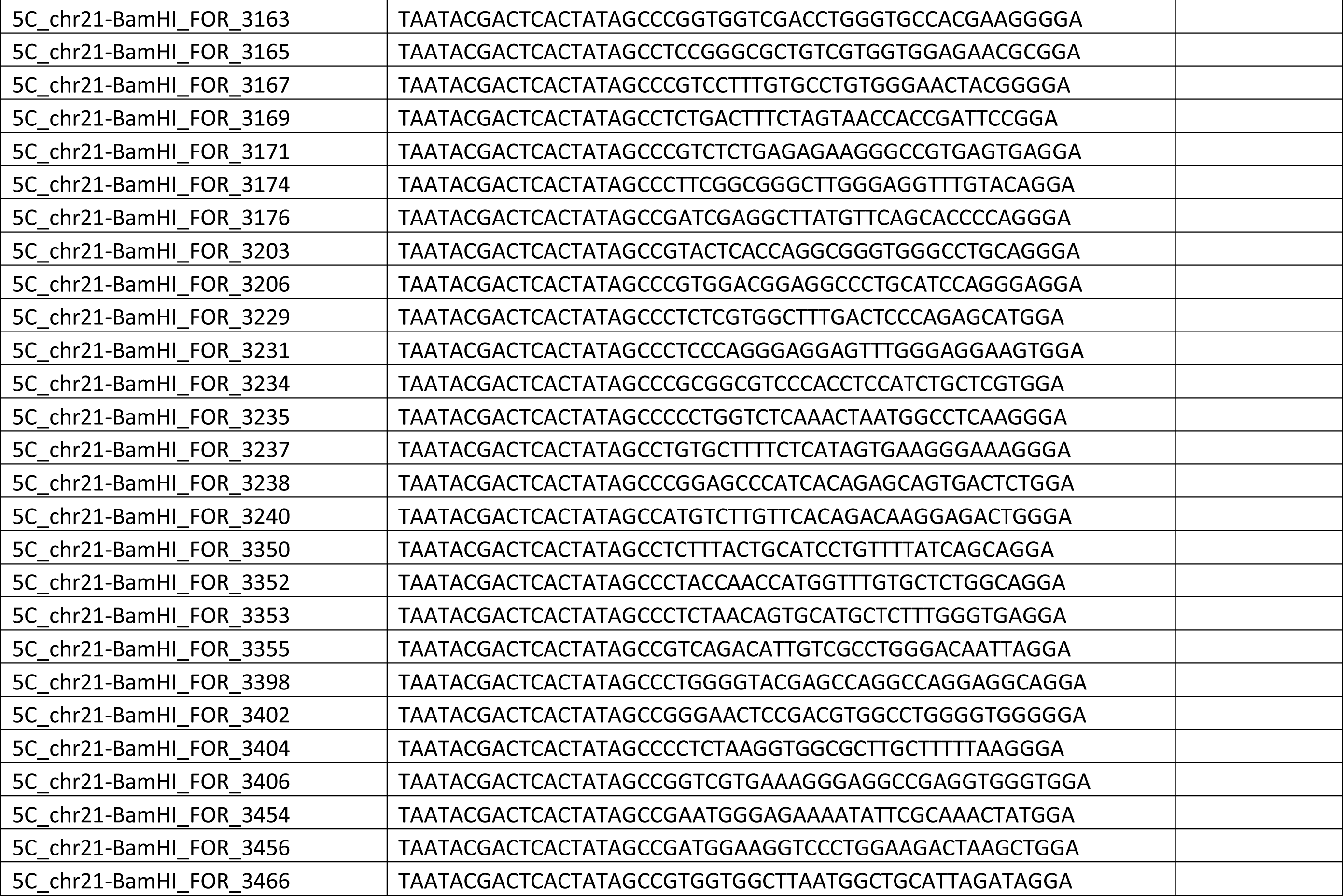
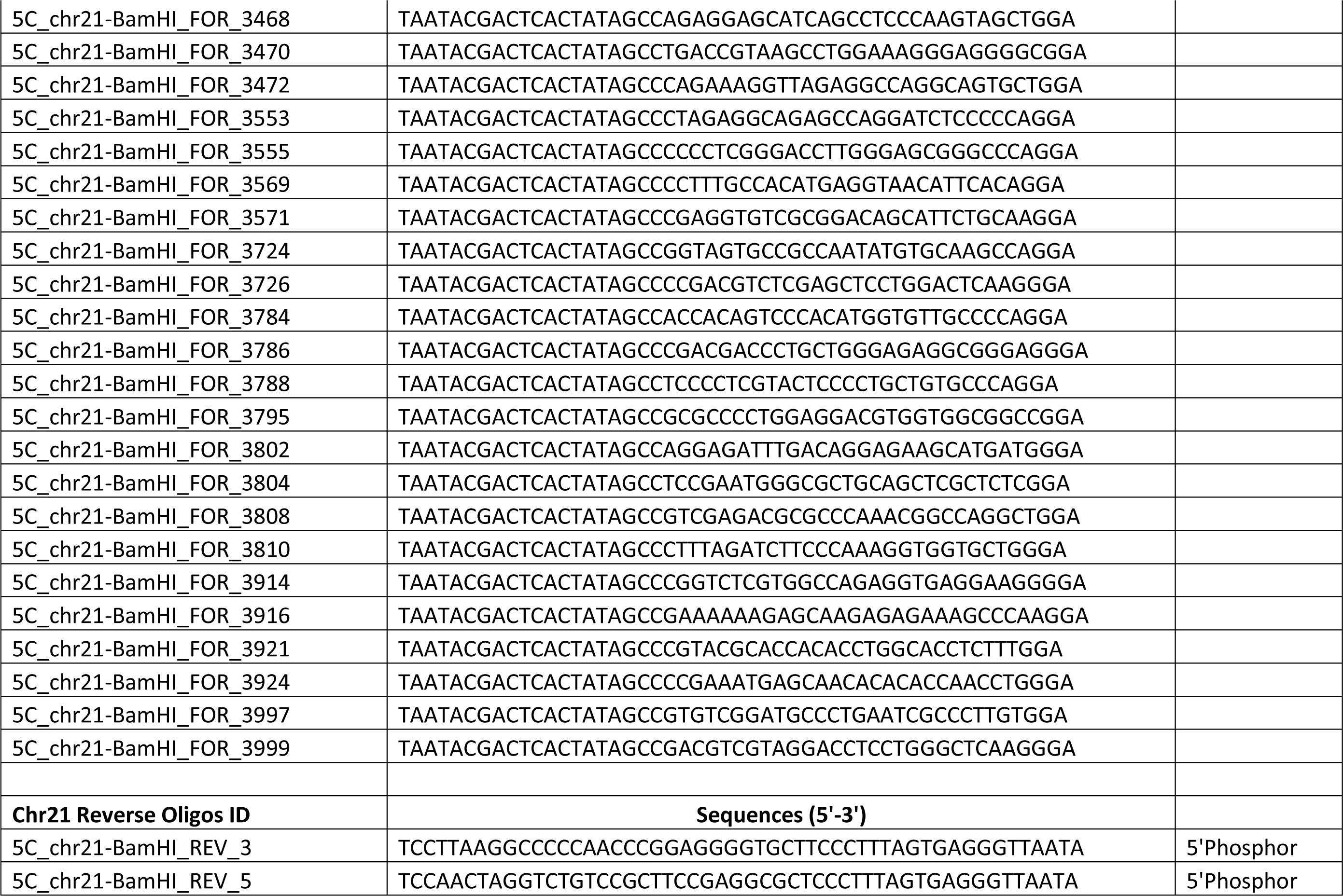
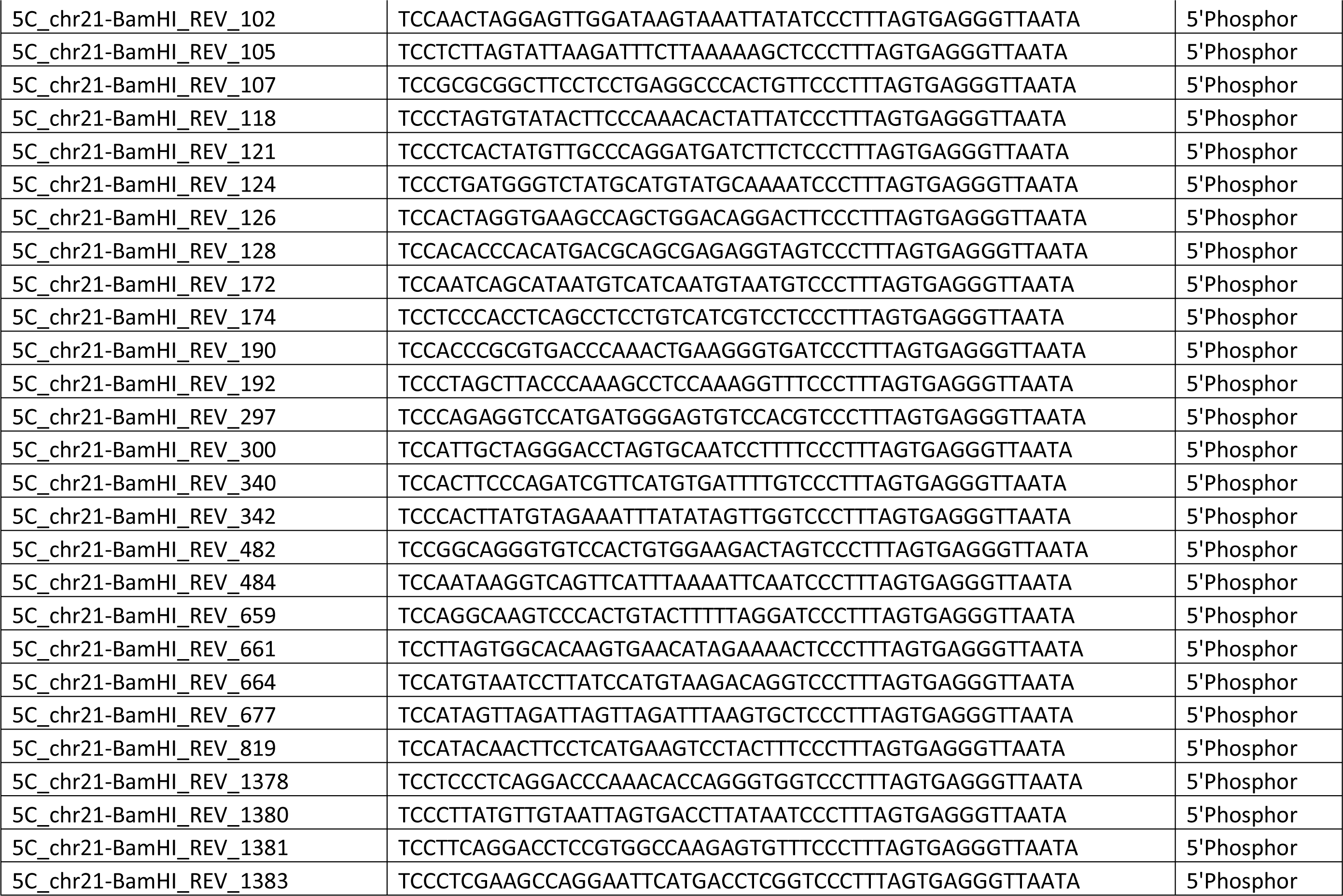
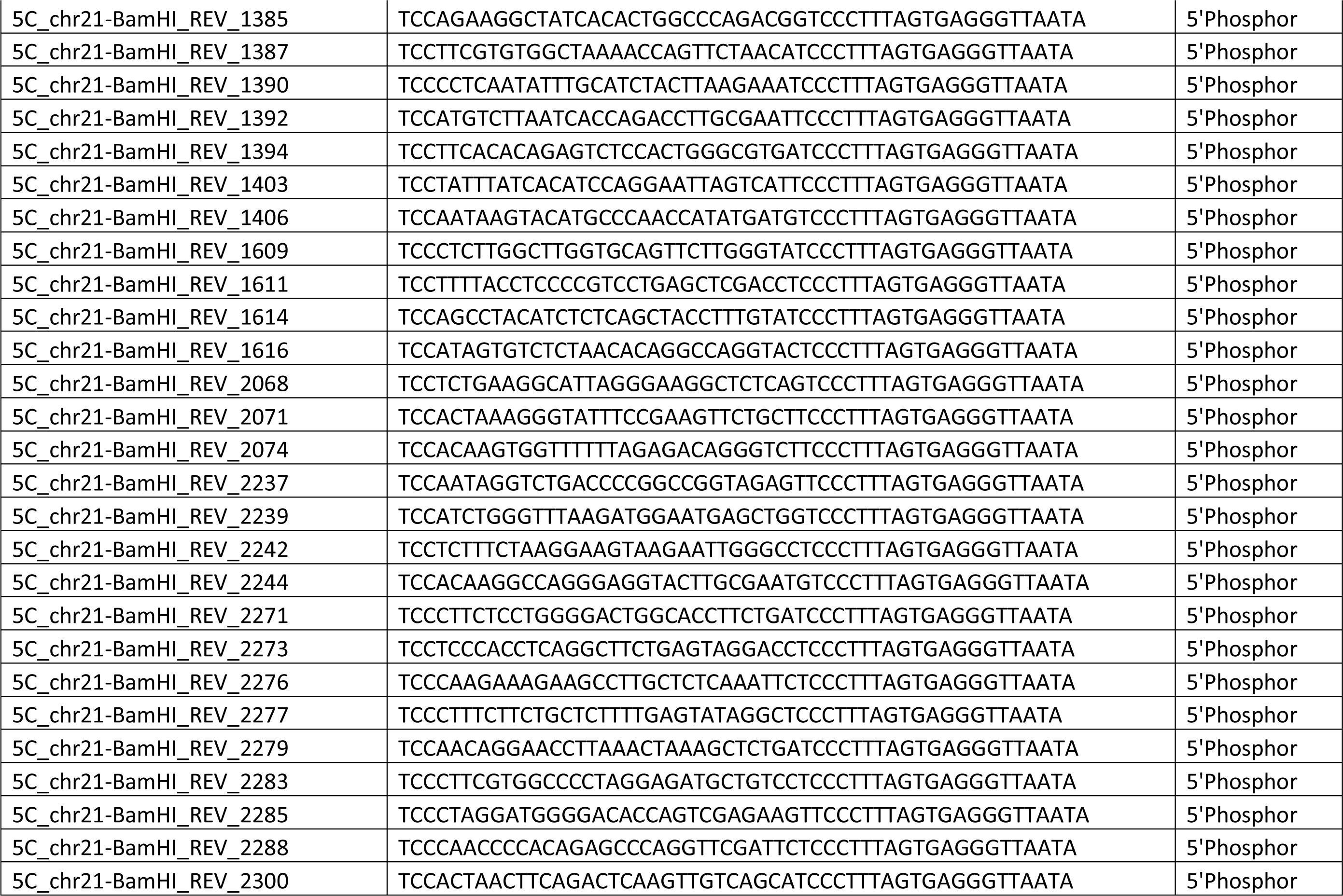
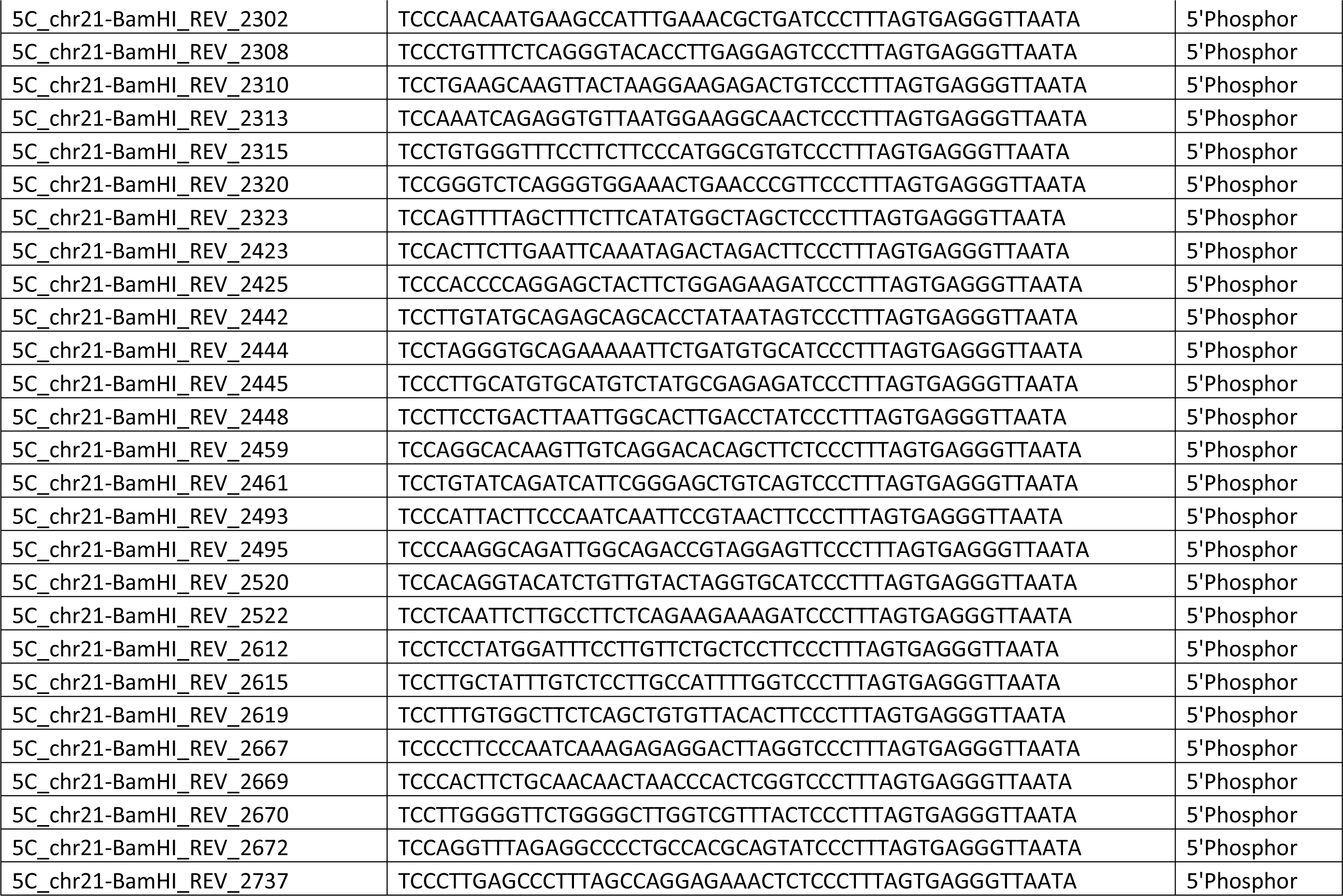
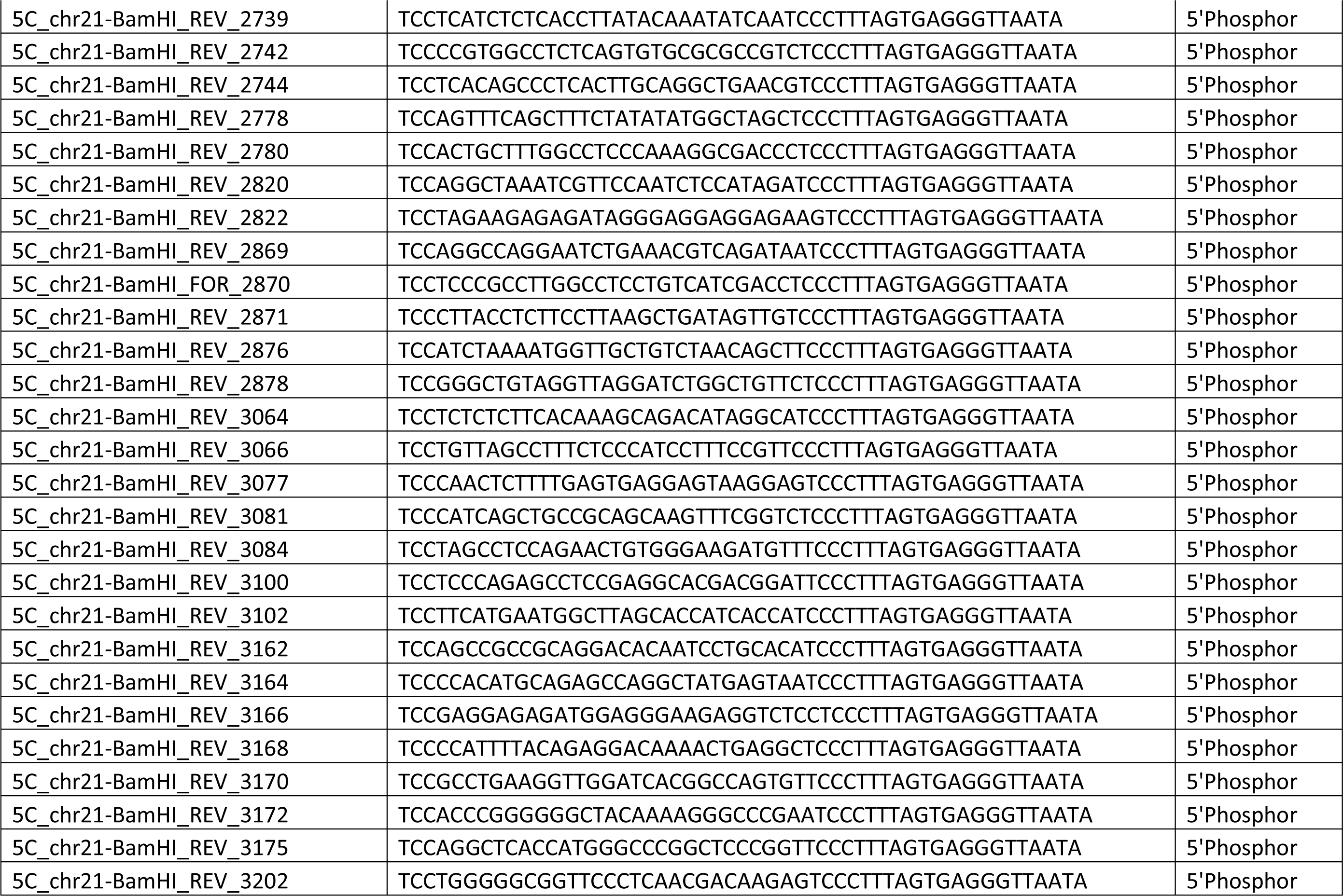
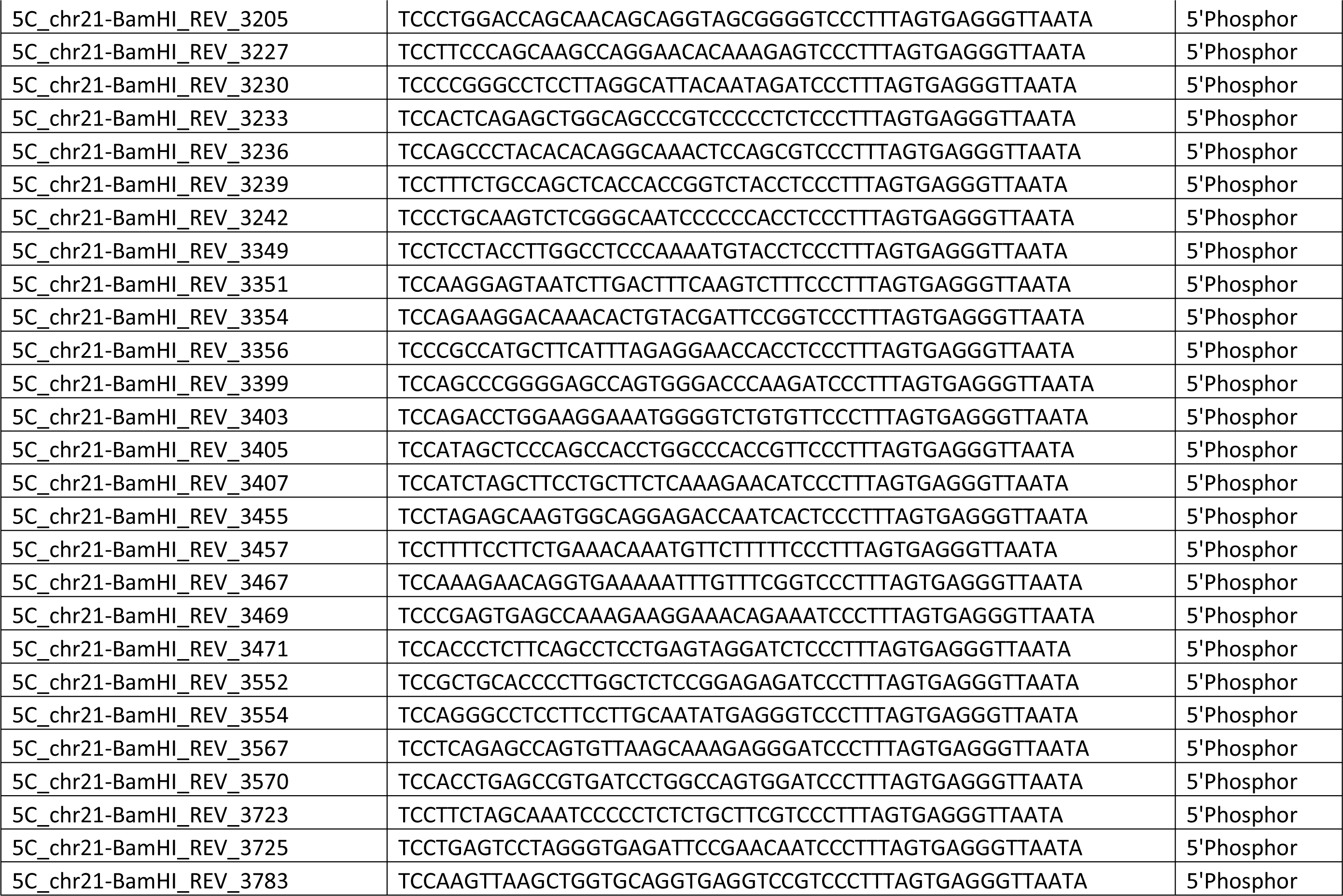
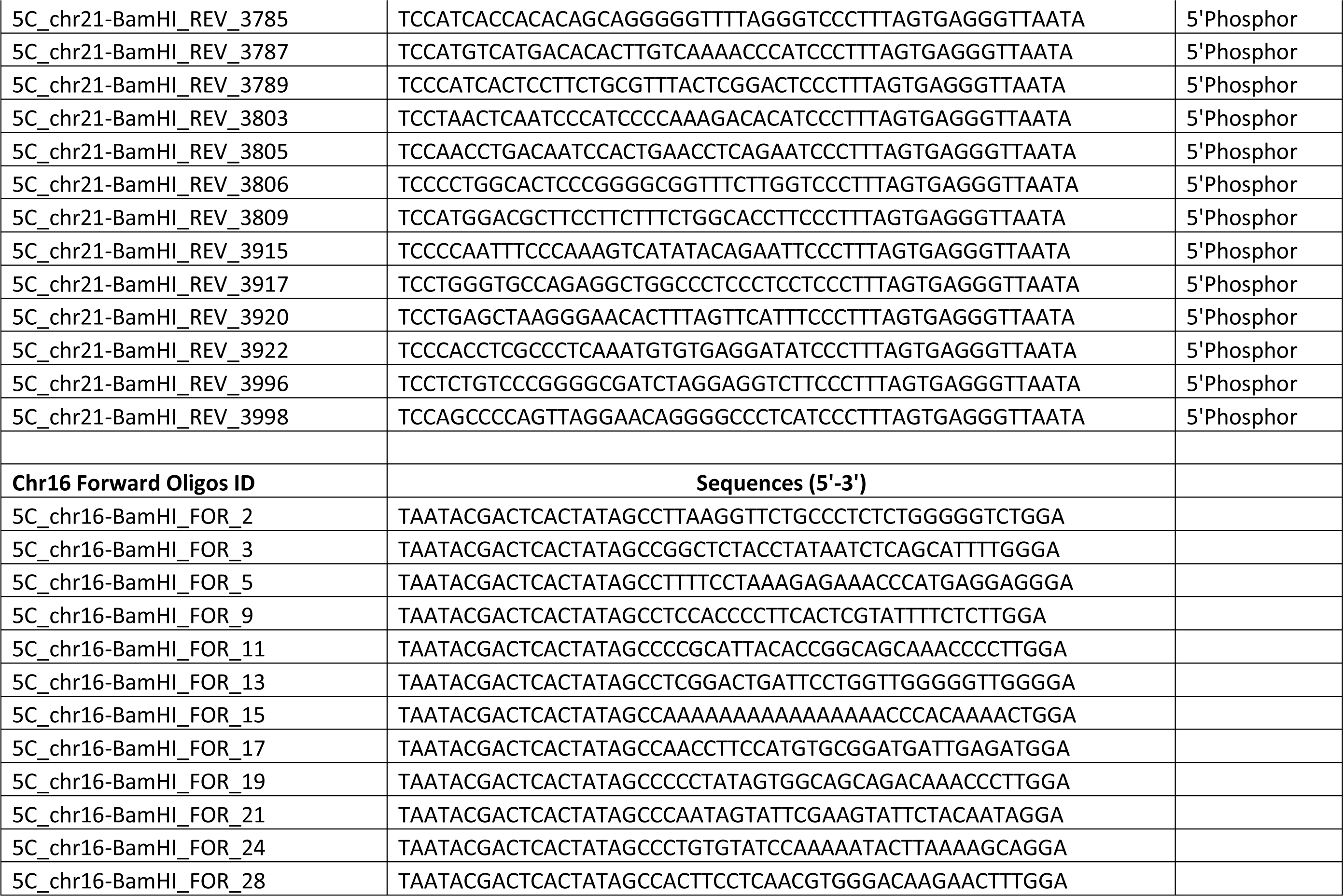
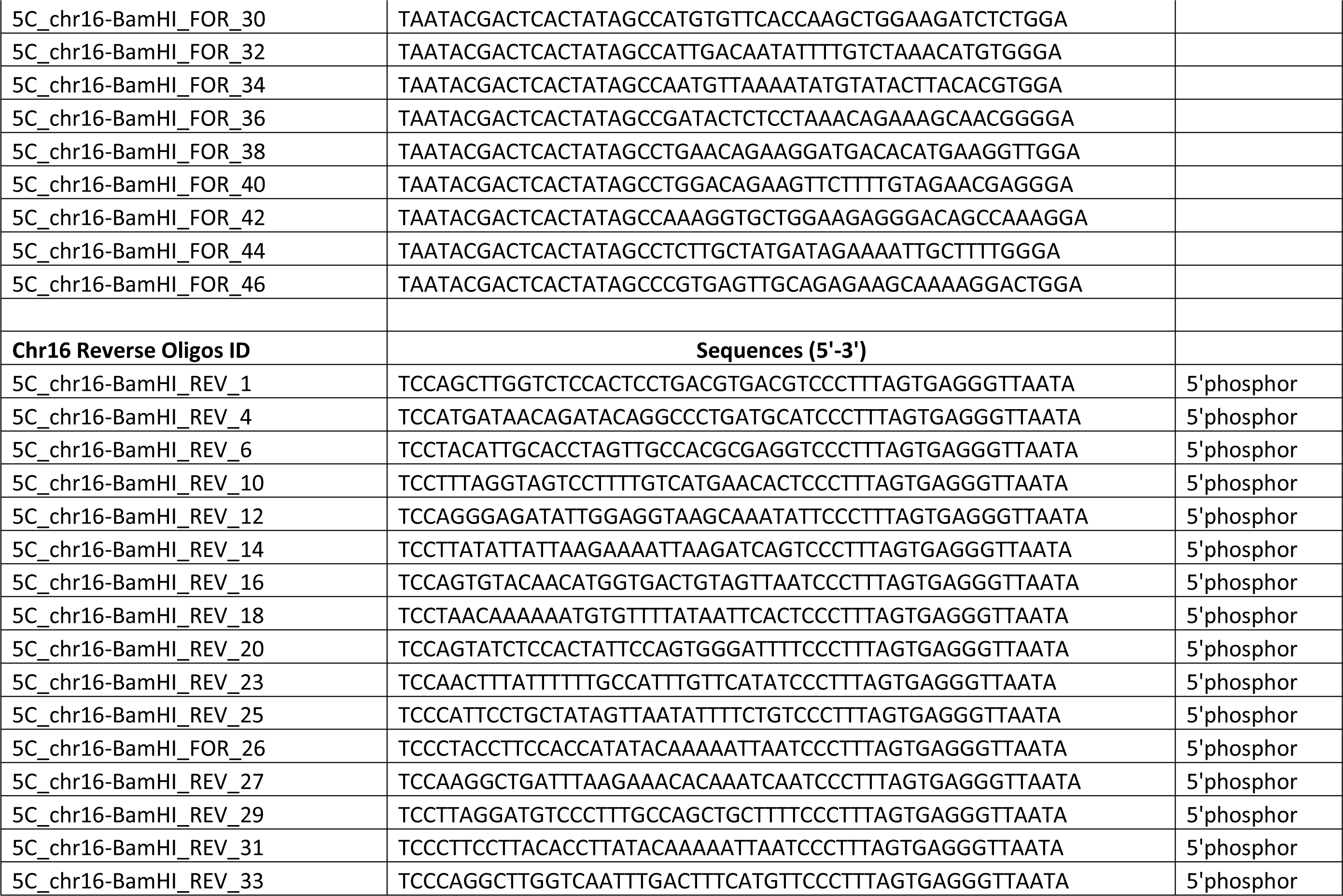
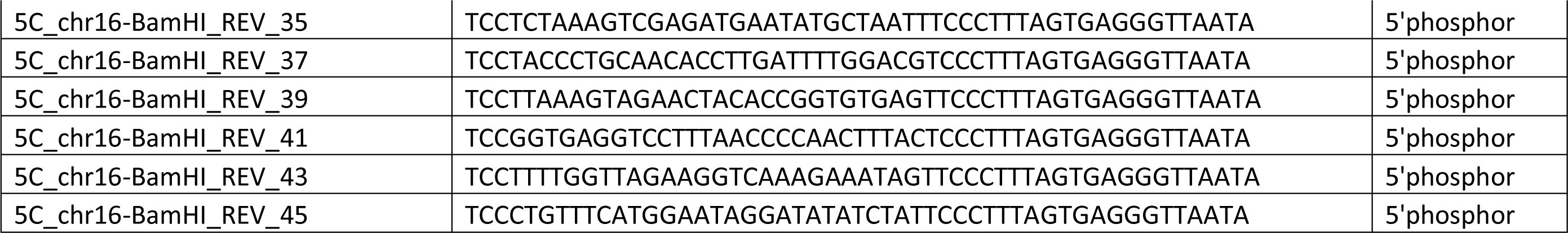
5C oligos used in the study

**Table S2:** 5C reads and Percent Alignment

| **Sample** | **Total Reads** | **Aligned Reads** | **Percent Alignment** | **Total Contacts** | **All Possible Contacts** |
| --- | --- | --- | --- | --- | --- |
| ICI 5C Rep1 | 25612149 | 25320119 | 0.988597989 | 17839 | 23095 |
| ICI 5C Rep2 | 25491622 | 25325094 | 0.993467344 | 17882 | 23095 |
| E2 5C Rep1 | 26623491 | 26140530 | 0.981859592 | 18206 | 23095 |
| E2 5C Rep2 | 26285590 | 26007082 | 0.989404537 | 18060 | 23095 |

**Table S3:**
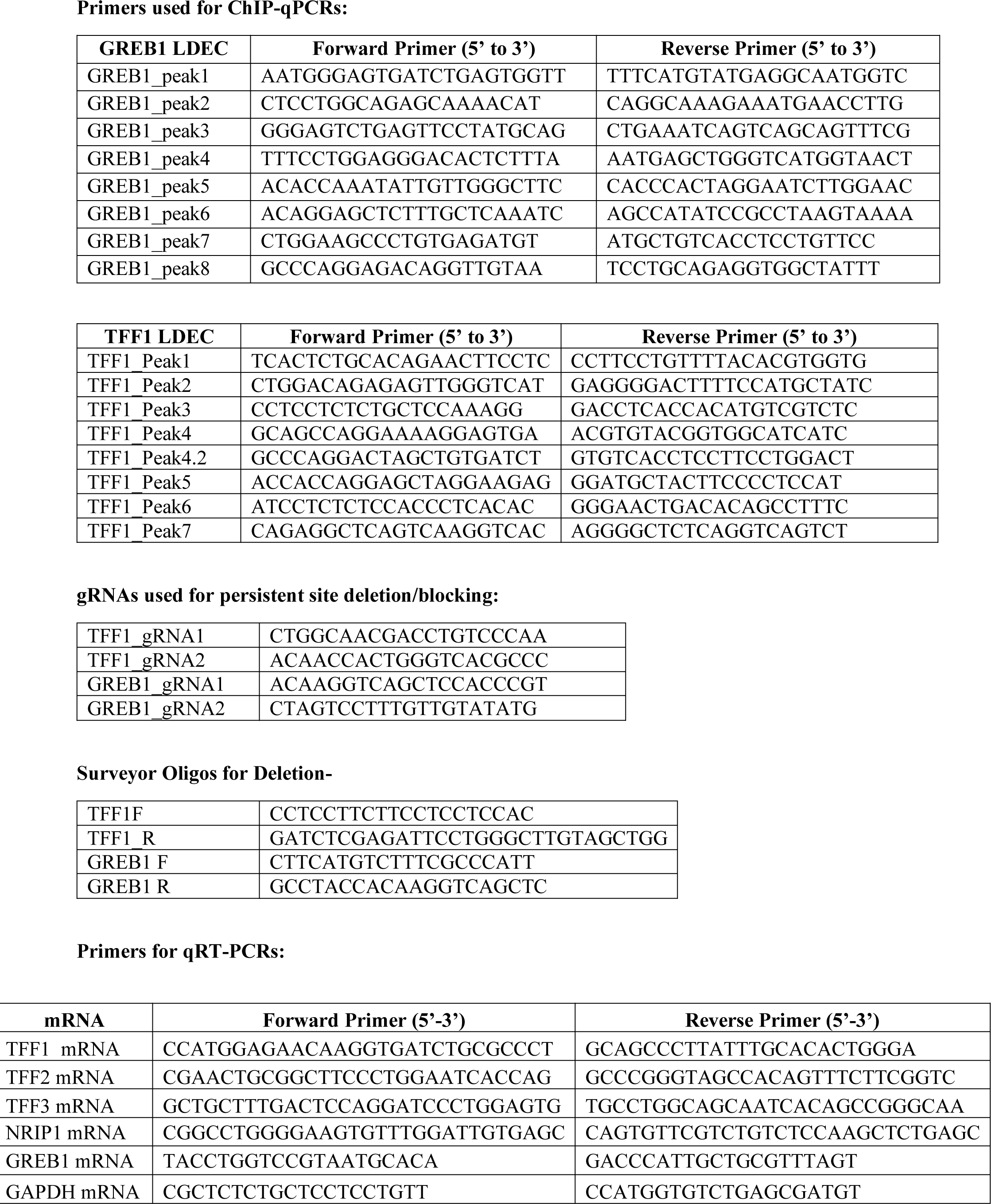
Sequences of gRNA and primers

**Table S4:**
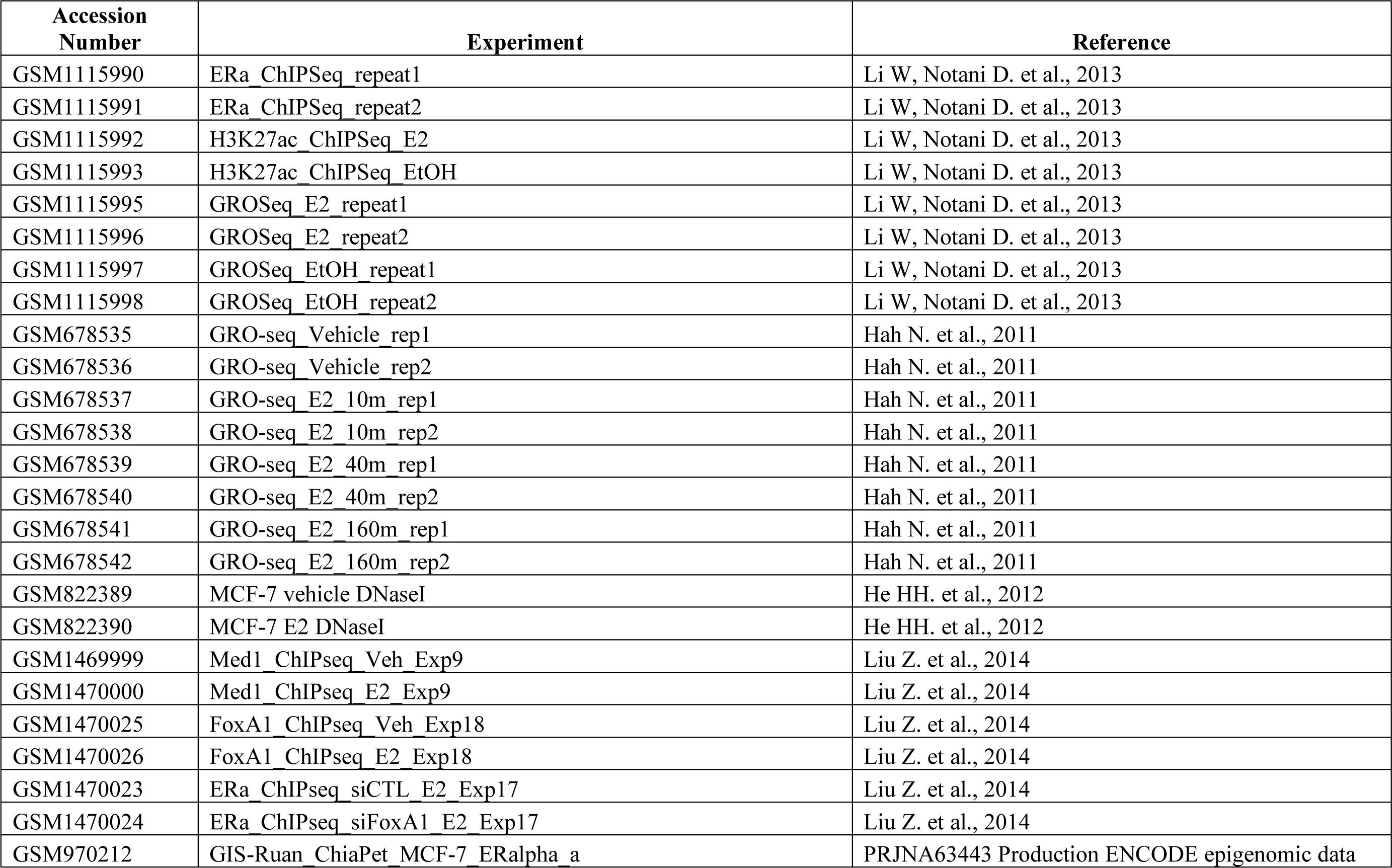
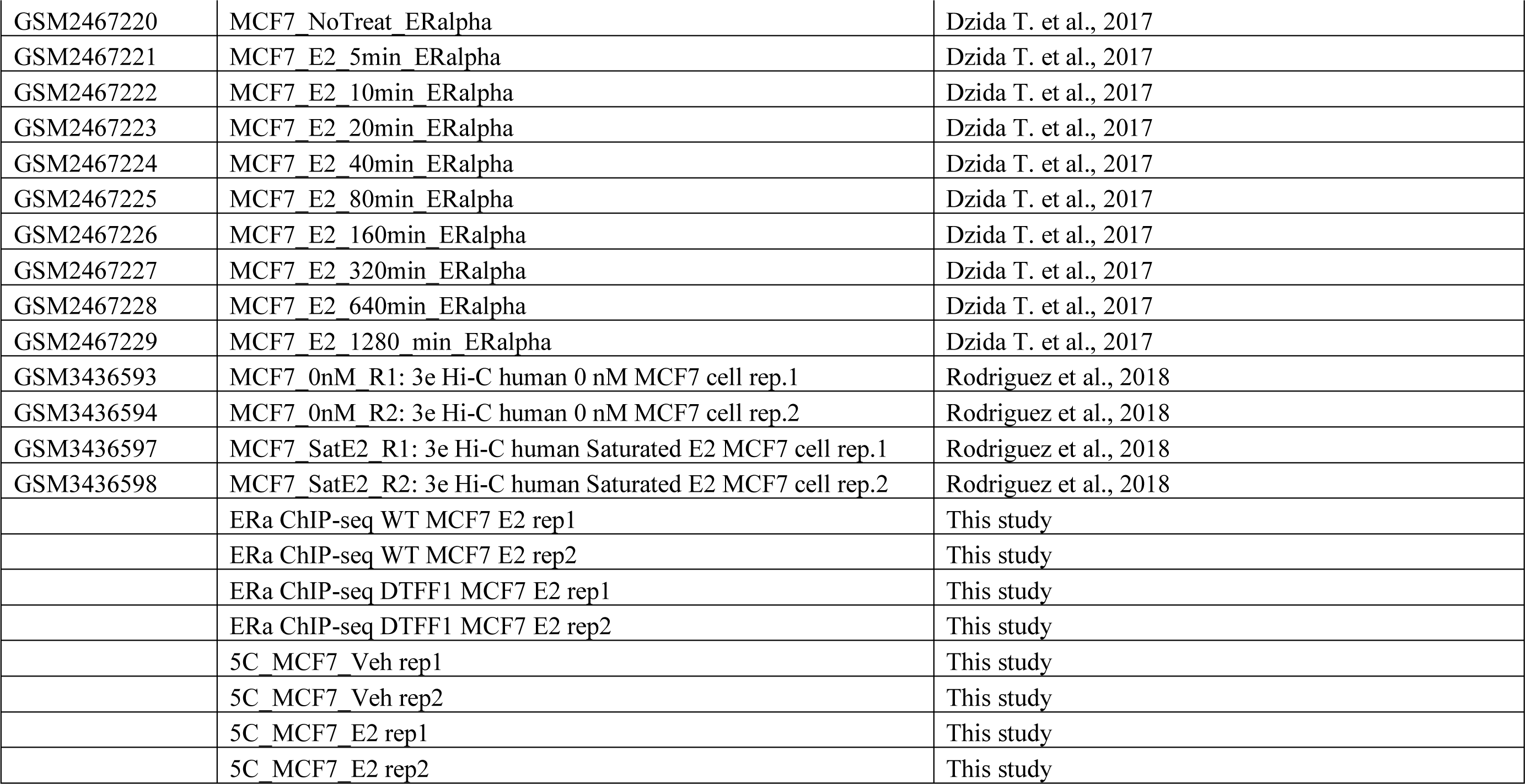
Accession Numbers of NGS datasets

